# Strongly deleterious mutations are a primary determinant of extinction risk due to inbreeding depression

**DOI:** 10.1101/678524

**Authors:** Christopher C. Kyriazis, Robert K. Wayne, Kirk E. Lohmueller

## Abstract

Human-driven habitat fragmentation and loss have led to a proliferation of small and isolated plant and animal populations with high risk of extinction. One of the main threats to extinction in these populations is inbreeding depression, which is primarily caused by the exposure of recessive deleterious mutations as homozygous by inbreeding. The typical approach for managing these populations is to maintain high genetic diversity, often by translocating individuals from large populations to initiate a ‘genetic rescue.’ However, the limitations of this approach have recently been highlighted by the demise of the gray wolf population on Isle Royale, which was driven to the brink of extinction soon after the arrival of a migrant from the large mainland wolf population. Here, we use a novel population genetic simulation framework to investigate the role of genetic diversity, deleterious variation, and demographic history in mediating extinction risk due to inbreeding depression in small populations. We show that, under realistic models of dominance, large populations harbor high levels of recessive strongly deleterious variation due to these mutations being hidden from selection in the heterozygous state. As a result, when large populations contract, they experience a substantially elevated risk of extinction after these strongly deleterious mutations are exposed by inbreeding. Moreover, we demonstrate that although translocating individuals to small populations is broadly effective as a means to reduce extinction risk, using small or moderate-sized source populations rather than large source populations can greatly increase the effectiveness of genetic rescue due to greater purging in these smaller populations. Our findings challenge the traditional conservation paradigm that focuses on maximizing genetic diversity to reduce extinction risk in favor of a view that emphasizes minimizing strongly deleterious variation. These insights have important implications for managing small and isolated populations in the increasingly fragmented landscape of the Anthropocene.

**Impact Summary:** Numerous threats to extinction exist for small populations, including the detrimental effects of inbreeding. Although much of the focus in reducing these harmful effects in small populations has been on maintaining high genetic diversity, here we use simulations to demonstrate that emphasis should instead be placed on minimizing strongly deleterious variation. More specifically, we show that historically-large populations with high levels of genetic diversity also harbor elevated levels of recessive strongly deleterious mutations hidden in the heterozygous state. Thus, when these populations contract, inbreeding can expose these strongly deleterious mutations as homozygous and lead to severe inbreeding depression and rapid extinction. Moreover, we demonstrate that, although translocating individuals to these small populations to perform a ‘genetic rescue’ is broadly beneficial, the effectiveness of this strategy can be greatly increased by targeting historically-smaller source populations where recessive strongly deleterious mutations have been purged. These results challenge long-standing views on how to best conserve small and isolated populations facing the threat of inbreeding depression, and have immediate implications for preserving biodiversity in the increasingly fragmented landscape of the Anthropocene.

## Introduction

The prevailing paradigm in conservation biology prioritizes the maintenance of high genetic diversity in small populations threatened with extinction due to inbreeding depression (Caughley 1994; Spielman et al. 2004). Under this paradigm, genetic diversity is considered one of the primary determinants of fitness (Allendorf and Leary 1986; Reed and Frankham 2003), and the harmful effects of inbreeding are thought to be minimized by maintaining genetic diversity. However, this thinking is challenged by the observation that some species, such as the Channel island fox, can persist at small population size with extremely low genetic diversity and show no signs of inbreeding depression (Robinson et al. 2016, 2018). This example and other similar studies suggest that, rather than being mediated by high genetic diversity, the persistence of small populations may instead be enabled by the purging of strongly deleterious mutations (Laws and Jamieson 2011; Xue et al. 2015; Hedrick and Garcia-Dorado 2016; Robinson et al. 2016, 2018; Van Der Valk et al. 2019; Grossen et al. 2020). In this study, we investigate how genetic diversity, deleterious variation, and demographic history influence extinction risk due to inbreeding depression using ecologically-realistic population genetic simulations. Our results demonstrate the central role of strongly deleterious variation in determining extinction risk due to inbreeding depression in small and isolated populations, and highlight the counterintuitive effects of strategies aimed at maintaining high genetic diversity. We argue that, in cases where populations are destined to remain small and isolated with high levels of inbreeding, management strategies should aim to minimize strongly deleterious variation rather than maximize genetic diversity.

The motivating example for our simulations is the gray wolf population on Isle Royale, an island in Lake Superior that has long served as a natural laboratory in ecology and conservation biology (Mech 1966; Peterson et al. 1984; Wayne et al. 1991; McLaren and Peterson 1994; Hedrick et al. 2019). Following 70 years of near-complete isolation at a size of ~10-50 individuals, the population was driven to the brink of extinction by severe inbreeding depression, with just two individuals remaining in 2018 (Hedrick et al. 2019; Robinson et al. 2019). Recent findings have suggested that the collapse of the population may have been prompted by the introduction of high levels of recessive strongly deleterious variation by a migrant individual who arrived from the mainland in 1997 (Adams et al. 2011; Hedrick et al. 2014, 2019; Robinson et al. 2019). The high reproductive output of this individual on the island (34 offspring) led to intensive subsequent inbreeding in the population, driving these strongly deleterious mutations to high frequency and leading to severe inbreeding depression and ultimately the collapse of the population.

This example, although extreme, highlighted the potentially negative effects of founding or rescuing small populations with individuals from large and genetically-diverse populations. An alternative approach for genetic rescue or reintroduction initiatives could instead target historically-smaller populations where strongly deleterious mutations have been purged, or screen populations for individuals with low levels of strongly deleterious variation. The growing evidence for purging in wild populations (e.g., Xue et al. 2015; Robinson et al. 2018; Grossen et al. 2020) suggests that this approach may be effective as a means to decrease the severity of inbreeding depression in small populations at high risk of extinction. Given the ongoing reintroduction of wolves to Isle Royale, and the increasing necessity of reintroduction and genetic rescue more broadly (Whiteley et al. 2015; Frankham et al. 2017; Bell et al. 2019), such an approach could have wide-ranging implications for the conservation of small populations at risk.

The applicability of population genetic models to understanding extinction has historically been limited by unrealistic assumptions that often ignore stochastic ecological factors and rarely consider both weakly and strongly deleterious mutations (Lande 1994; Lynch et al. 1995; O’Grady et al. 2006; Caballero et al. 2017). Here, we use a novel population genetic simulation framework that combines realistic models of population dynamics with exome-scale genetic variation (Haller and Messer 2019) to assess how genetic diversity, deleterious variation, and demographic history influence extinction risk in small populations. Our simulations aim to capture the ecological factors that may contribute to extinction in small populations, such as those observed in the Isle Royale wolf population, by incorporating the effects of demographic and environmental stochasticity, as well as natural catastrophes. Coupled with these stochastic population dynamics, we model a genome with parameters reflecting that of a canine exome, which accumulates neutral and deleterious mutations from an empirically-estimated distribution of fitness effects (Kim et al. 2017). Although our model is motivated by the Isle Royale wolf population, it is also intended to capture the dynamics of many other classic examples of population decline, inbreeding depression, and genetic rescue, such as Scandinavian wolves (Åkesson et al. 2016), Florida panthers (Johnson et al. 2010), and bighorn sheep (Hogg et al. 2006).

## METHODS

### Overview of the SLiM non-Wright-Fisher model

We conducted non-Wright-Fisher (nonWF) simulations using SLiM 3 (Haller and Messer 2019). The impetus for this model was to allow for more ecologically-realistic population genetic simulations by relaxing many of the restrictive assumptions of the Wright-Fisher model (Haller and Messer 2019). Such assumptions include non-overlapping generations and a fixed population size that is not influenced by fitness, both of which are unrealistic when trying to model the extinction of a population due to genetic factors.

Instead, the SLiM nonWF approach models population size (N) as an emergent property of individual absolute (rather than relative) fitness and a user-defined carrying capacity (K). Thus, if individual fitness declines, a population experiences extinction through a biologically realistic process (a fitness-driven reduction in population size). Further, the model includes overlapping generations, such that individuals with high fitness can survive and reproduce for multiple generations. At the start of each generation, each individual randomly mates with another individual in the population, with one offspring resulting from each mating. At the end of each generation, individuals die with a probability given by their absolute fitness (ranging from 0 to 1), which is rescaled by the ratio of K/N to model the effects of density dependence. Thus, the carrying capacity here does not directly determine the simulated population size, but rather it indirectly influences it through its impact on individual fitness and viability selection. For the sake of both tractability and generality, we assume a hermaphroditic random mating population. A discussion of how the carrying capacity of a SLiM nonWF model is related to its Wright-Fisher effective population size is provided in the Supplementary Methods.

### Demographic scenarios

To obtain a baseline understanding for how ancestral demography influences extinction risk in small populations, we first explored a ‘population contraction’ scenario with our simulations (Figure 1A). Here, we modeled an instantaneous contraction from a large ‘ancestral population’ (K_ancestral_∈ {1,000, 5,000, 10,000, 15,000}) to a small ‘endangered population’ (K_endangered_ ∈ {25, 50}) (Figure 1A). This contraction scenario could represent the isolation of a population with historical connectivity (e.g., the Florida panther population) or alternatively the founding of an isolated population through migration or translocation (e.g., the initial founding of the Isle Royale wolf population). For each contraction event, we randomly sampled K_endangered_ number of individuals from the ancestral population to seed the endangered population after a burn in of 10*K_ancestral_ generations. All simulations were run until the endangered population went extinct. To examine the effects of a more gradual contraction, we also explored a scenario where an ancestral population with carrying capacity 10,000 first contracted to an intermediate carrying capacity of 1,000 for 200 generations, and finally an endangered carrying capacity of 25. We ran 25 simulation replicates for each combination of ancestral and endangered carrying capacities.

**Figure 1:**
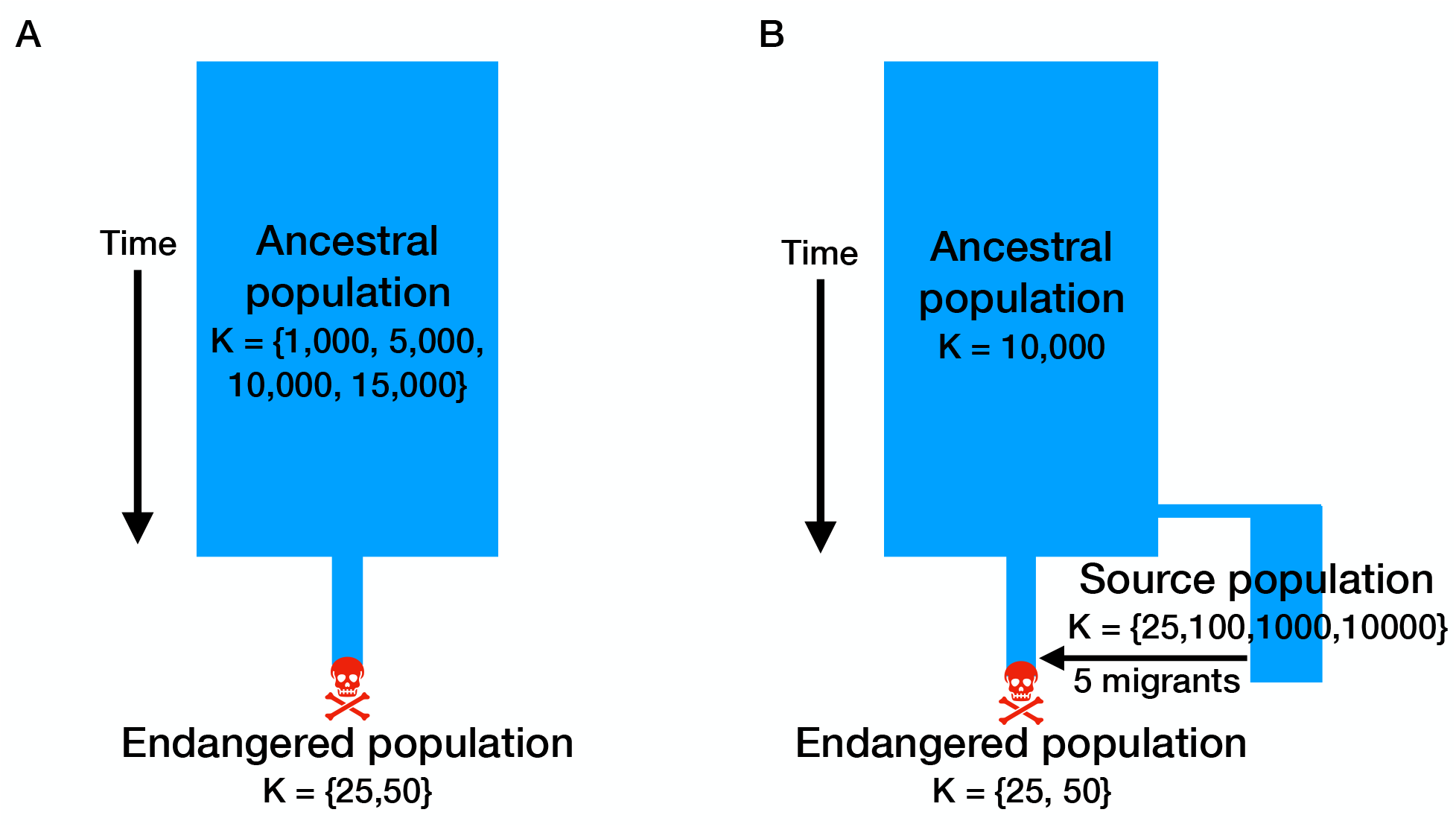
Schematic of the demographic scenarios used for simulations. (A) Schematic for ‘population contraction’ simulations. (B) Schematic for ‘genetic rescue’ simulations. Note that migration occurs in the genetic rescue scenario when the endangered population decreases in size to five or fewer individuals when K_endangered_ =25 and 15 or fewer individuals when K_endangered_ =50.

We next explored a ‘genetic rescue’ scenario, which similarly consisted of a population contraction from a large ancestral population to a small endangered population (Figure 1B). Here, however, we fixed the ancestral carrying capacity to 10,000, and again explored two endangered carrying capacities of 25 and 50. Prior to the contraction, we split off the following source populations for genetic rescue: 1) a large population remaining at the ancestral size (K=10,000); 2) a moderate-sized population with long-term isolation (K=1,000 for 1,000 generations); 3) a small population with relatively recent isolation (K=100 for 100 generations); and 4) a very small population with very recent isolation (K=25 for 10 generations). These source population demographic histories were set to reflect a range of biologically-relevant scenarios (i.e., large and outbred populations, populations with moderate size and long-term isolation, smaller populations that have been more recently isolated) as well as provide a range of source population levels of genetic diversity and strongly deleterious variation to examine how these factors influence the efficacy of genetic rescue. Genetic rescue was initiated by translocating five randomly-selected individuals after the endangered population decreased in size to five or fewer individuals for the case when K_endangered_=25 and 15 or fewer individuals for the case when K_endangered_=50. Importantly, the exact number of generations of isolation for these source populations depended on the number of generations before translocation, which varied for each simulation replicate depending on the stochastic trajectory of the endangered population. For these simulations, we ran 50 replicates for simulations with K_endangered_=25 and 25 replicates for simulations with K_endangered_=50.

To further explore the dynamics of our genetic rescue scenario, we ran several additional simulations, here with endangered carrying capacity fixed to 25. First, to investigate the impact of selecting individuals for translocation to either maximize genetic diversity or minimize strongly deleterious variation, we ran simulations (50 replicates) were we picked selected five migrants from the K=10,000 source population with either the highest heterozygosity or fewest number of strongly deleterious alleles (where strongly deleterious mutations were defined to be those mutations with *s* < −0.01). We also explored scenarios with varying numbers of migrants (1, 5, or 10) with the number of translocations fixed to one, as well as varying the number of translocations (1, 2, or 5) with the number of migrants fixed to five. Here, we ran 25 replicates for each parameter combination. All simulations were run until the endangered population went extinct.

### Stochastic population dynamics

To capture the non-genetic factors that can contribute to extinction in small populations (Caughley 1994), our model includes three sources of ecological stochasticity. First, demographic stochasticity was modelled using the built-in mechanics of the SLiM nonWF model, in which survival from one generation to the next is determined by a Bernoulli trial with the probability of survival determined by the absolute fitness of an individual scaled by K/N (Haller and Messer 2019). Next, we incorporated the effects of environmental stochasticity in our simulations by modelling the carrying capacity of the endangered population as an Ornstein-Uhlenbeck process, in which the carrying capacity in a generation is given by:

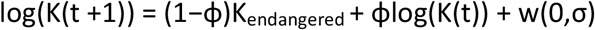

where ϕ = 0.9, K_endangered_ ∈{25,50} and σ = log_10_(1.3) (Figure S1). The values of ϕ and σ were set arbitrarily to model environmental stochasticity with a moderate amount of variation and a high degree of auto-correlation. Finally, we modelled the effects of random natural catastrophes in our simulations by drawing a probability of mortality due to a catastrophe each generation from a beta distribution with α= 0.5 and β= 8 (Figure S1). In each generation, deaths due to a catastrophe are then determined by the outcome of a Bernoulli trial for each individual with the probability given by the beta distribution. Environmental stochasticity and natural catastrophes were only modelled in the small endangered population. Importantly, these stochastic ecological effects rarely lead to extinction in the endangered population in the absence of deleterious variation (8/25 simulation replicates with neutral mutations extinct within 10,000 generations at K_endangered_ = 25, and none extinct for K_endangered_ = 50).

### Genomic parameters

We set the genomic parameters in our simulations to model the exome of a wolf-like organism. To do this, each diploid individual in our simulation has 20,000 genes of length 1500 bp, which occur on 38 autosomes with the number of genes on each chromosome determined by the ratios observed in the dog genome (Lindblad-Toh et al. 2005). Recombination between these genes occurs at a rate of 1 × 10^−3^, with no recombination within genes and free recombination between chromosomes. These genes accumulate neutral and deleterious mutations at a rate of 1 × 10^−8^ per site, with the ratio of deleterious to neutral mutations set to 2.31:1 (Huber et al. 2017; Kim et al. 2017). The selection coefficients for these deleterious mutations were drawn from a distribution of fitness effects (DFE) estimated from a large sample of humans (Kim et al. 2017; see Supplementary Methods for additional details).

We initially set the dominance coefficients for our simulations according to a *hs* relationship inferred for yeast (Agrawal and Whitlock 2011) following the approach from Henn et al. 2016 with the following functional form:

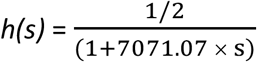

This *hs* relationship intends to capture the pattern evident in empirical data that the dominance coefficient of a mutation tends to be inversely related to its selection coefficient, such that the most strongly deleterious mutations are highly recessive (Simmons and Crow 1977; Agrawal and Whitlock 2011; Huber et al. 2018). However, we found that simulations with this *hs* relationship in SLiM were extremely computationally intensive (see Supplementary Methods), such that we were only able to obtain results for simulations with the smallest ancestral carrying capacities of 1,000 and 5,000.

To overcome these computational limitations for realistic models of dominance, we instead implemented an approach assuming that weakly/moderately deleterious mutations (*s* ≥ −0.01) were partially recessive (*h*=0.25) and strongly deleterious mutations (*s* < −0.01) were fully recessive (*h*=0). The aim of this approach (hereafter referred to as our “*hmix”* model) was to capture the key feature of the *hs* relationship that more deleterious mutations tend to be more recessive, with the dominance coefficients for these two classes of mutations reflecting their mean dominance coefficient under the *hs* relationship (Figure S2). Given our finding that the behavior of this model is extremely similar to that of the *hs* relationship (see Results), we use the *hmix* model for all simulations except where otherwise noted. More details on how the *hmix* model was implemented in SLiM is provided in the Supplementary Methods.

To further explore the impact of dominance coefficients, we also ran simulations where we varied the dominance coefficient for all mutations (*h* ∈ {0, 0.01, 0.05, 0.2, 0.5}). In addition, we explored the impact of decreasing the number of sites in which fully recessive (*h*=0) deleterious mutations can occur (i.e. the mutational target size) by varying the number of genes in our simulations (*g* ∈ {1,000, 5,000, 10,000, 15,000, 20,000}). These simulations were run solely under the ‘population contraction’ demographic scenario with K_ancestral_ ∈{1,000, 5,000, 10,000, 15,000} and K_endangered_ = 25. We ran 25 replicates for each of these simulations, terminating the simulation when the endangered population went extinct.

During the simulations, we kept track of several summaries of the state of the population. These include the population’s mean heterozygosity, mean inbreeding coefficient (F_ROH_), the mean fitness, and the average number of deleterious alleles per individual binned into weakly (−0.001 < *s* ≤ −0.00001), moderately (−0.01 < *s* ≤ −0.001), strongly (*s* < −0.01), and very strongly (*s* < −0.05) deleterious classes. Fitness was calculated multiplicatively across sites. Here, we restricted F_ROH_ to include only runs of homozygosity greater than 1 Mb to capture inbreeding due to mating between close relatives. These statistics were calculated from a sample of 30 individuals every 1000 generations during the burn-in and every generation following the contraction.

### Burn-in conditions for the simulations

Our simulations during the burn-in retained fixed mutations and did not model reverse mutation. Retaining fixed mutations during the burn-in was important to ensure that weakly deleterious mutations that drifted to fixation contribute to absolute fitness. However, one consequence of retaining fixed mutations is that there is no mutation-selection-drift equilibrium in the ancestral population since weakly deleterious mutations continue to accumulate as fixations even after heterozygosity of segregating variation has leveled off (Figure S3). As a result, fitness during the burn-in also never reaches equilibrium, but instead declines gradually as weakly deleterious mutations are fixed. Although fixation of weakly deleterious mutations occurs at a rate that is inverse to population size (i.e., much faster in smaller populations), we found that this effect is cancelled out when the length of the burn-in is proportional to the population’s carrying capacity (i.e., 10*K, leading to a much shorter burn-in for smaller populations), resulting in the same pre-contraction fitness regardless of population size (Figure S4a). To explore the effect of reducing the pre-contraction fitness of the smaller populations, we ran simulations under a fully recessive model (*h*=0) where we added an additional 20,000 generations to the burn-in for the K=1,000, K=5,000, and K=10,000 populations, leading to burn-in durations of 30,000, 70,000, and 120,000 respectively (10*K + 20,000). This led to a notable reduction in pre-contraction fitness for the K=1,000 population and more moderate reductions in fitness for the larger populations (Figure S4). However, there was no differences in time to extinction (Figure S4C), suggesting that extinction dynamics are governed more by the numbers of recessive strongly deleterious alleles in these populations rather than their pre-bottleneck fitness. This finding is further supported by our simulation results that demonstrate that strongly deleterious mutations are a far more important determinant of extinction times compared to weakly or moderately deleterious mutations (see Results). Thus, for the sake of computational tractability, we ran all burn-ins for 10*K_ancestral_ generations.

## Results

### Population contraction simulations under *hmix* model

Our population contraction simulations demonstrate that, although larger populations have higher genetic diversity (Figure 2A), they also harbor higher levels of strongly deleterious variation (Figure 2B). Consequently, we observe a strong effect of ancestral demography on time to extinction following a population contraction, with populations that were historically large experiencing more rapid extinction (Figure 2C). For example, given an endangered carrying capacity of 25, a population with an ancestral carrying capacity of 1,000 will go extinct in 474 generations on average, whereas a population with an ancestral carrying capacity of 15,000 will do so in 70 generations on average (Figure 2C). When modelling a more gradual contraction from an ancestral carrying capacity of 10,000, we found that extinction times were slightly increased relative to the instantaneous contraction scenario (Figure S5), suggesting that more gradual contractions can facilitate purging and ultimately decrease extinction risk.

**Figure 2:**
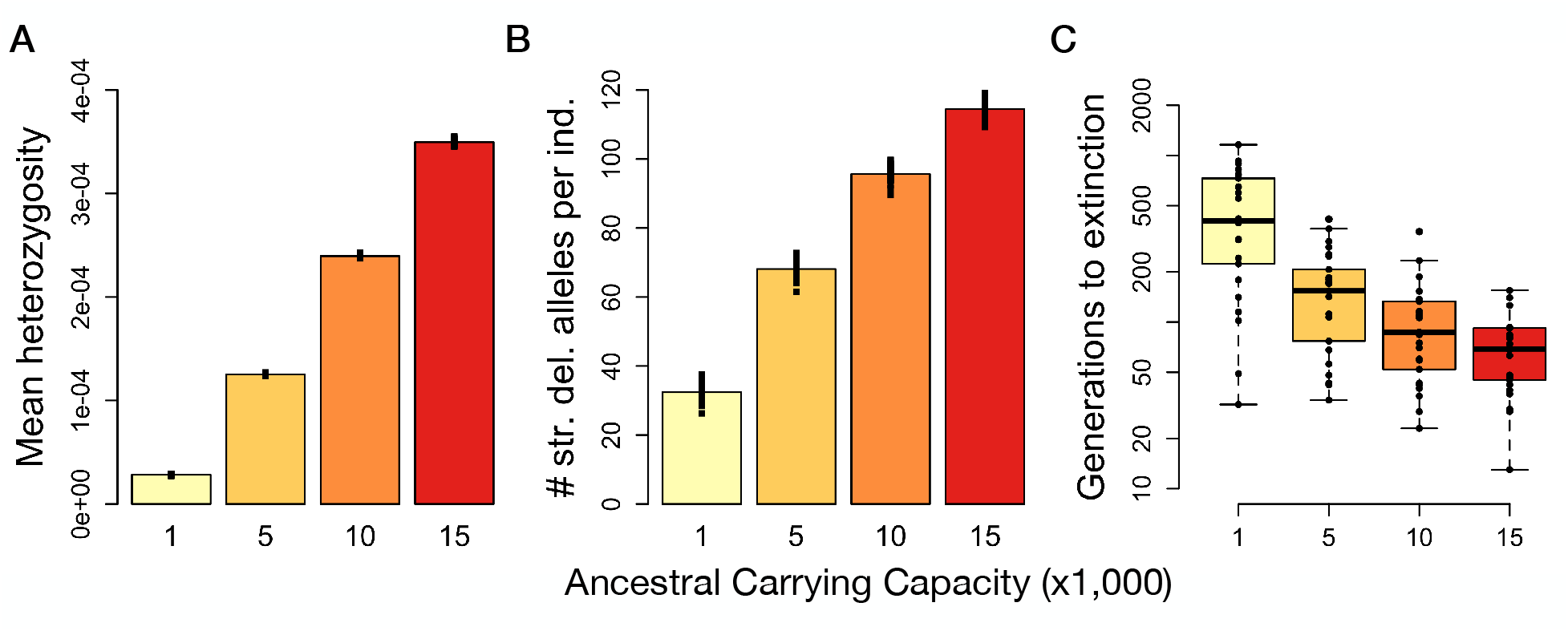
Results for population contraction simulations under the *hmix* model of dominance. (A) Mean heterozygosity of ancestral populations prior to contraction. (B) Average number of strongly deleterious alleles (*s* < −0.01) per individual in the ancestral populations prior to contraction. (C) Time to extinction following contraction from ancestral populations of varying size to an endangered population with K=25. Extinction times for K_endangered_ = 50 shown in Figure S8. Note that the y-axis is on a log scale. For each parameter setting, 25 simulation replicates were run.

Examination of individual simulation replicates provides insight into the dynamics of extinction in these populations (Figures 3A and S6). Endangered populations with an ancestral carrying capacity of 1,000 contain fewer strongly deleterious mutations, translating to a decreased severity of inbreeding depression and ultimately longer persistence (Figures 3A and S6). This decreased severity of inbreeding depression allows these populations to become highly inbred (F_ROH_ ≈ 1) well before going extinct (Figures 3A and S6). By contrast, endangered populations with a larger ancestral carrying capacity of 15,000 have much higher levels of recessive strongly deleterious variation due to these mutations being hidden from selection in the ancestral population, resulting in a more rapid loss of fitness as these populations become inbred (Figures 3B and S6). As a result, these populations typically go extinct well before F_ROH_ approaches one (Figures 3B and S6). These populations also lose fitness due to increased genetic drift and inbreeding among more distant relatives, which is not captured by our definition of F_ROH_.

**Figure 3:**
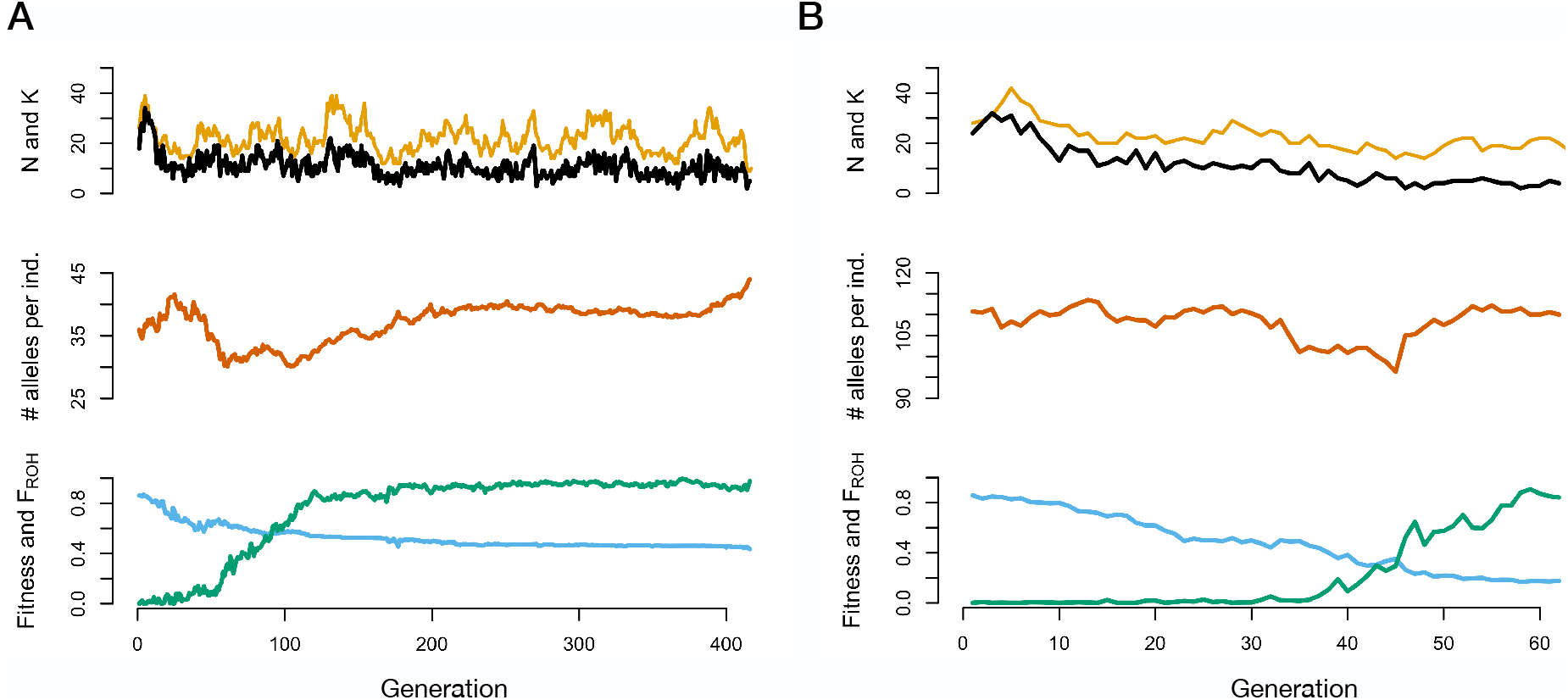
Representative examples of population contraction simulations with ancestral populations of varying size. (A) Example with K_ancestral_ = 1,000. (B) Example with K_ancestral_ = 15,000. The top of each panel shows population size (N) in black and carrying capacity (K) in gold, the middle shows the average number of strongly deleterious alleles (*s* < −0.01) per individual, and the bottom shows mean absolute fitness in blue and mean inbreeding coefficient (F_ROH_) in green. For both replicates, K_endangered_ = 25 and an *hmix* model of dominance was assumed. Note the differing y-axis scales for the middle panels and differing x-axis scales for panels A and B.

Following contraction, the ability of the endangered population to purge its load of recessive deleterious mutations also depended on stochastic ecological factors. For example, when the carrying capacity of the endangered population was, by chance, higher due to environmental stochasticity, natural selection was most effective at removing strongly deleterious alleles, translating to longer persistence (Figures 3 and S6). By contrast, when the carrying capacity of the endangered population was low soon after contraction, purging tended to be less effective, resulting in more rapid extinction (Figures 3 and S6). However, in both cases, purging was also counteracted by continual input of strongly deleterious alleles by mutation, which eventually contributed to population extinction. Overall, these various sources of genetic and ecological stochasticity together resulted in highly variable extinction times for any given parameter settings, highlighting the important role of random events in determining whether a small population can persist.

Our simulations also demonstrate the importance of increasing the carrying capacity of the endangered population as a means to ensure population persistence. Larger endangered populations (K=50) were better able to purge recessive deleterious mutations following the contraction, resulting in much longer persistence (Figure S7). Moreover, larger populations were less impacted by stochastic ecological factors, which also contributed to increased time to extinction. Nevertheless, extinction times for these larger endangered populations still depended strongly on the ancestral carrying capacity (Figure S7), demonstrating the importance of considering both recent and historical demography when assessing extinction risk.

### Results under varying genomic parameters

We next explored the sensitivity of our results to the genomic parameters assumed in our model. In particular, we focus on the impact of dominance coefficients, as previous work has suggested that the extent to which strongly deleterious mutations accumulate at higher frequencies in larger populations depends strongly on the assumed dominance coefficients (Nei 1968; Hedrick 2002; Hedrick and Garcia-Dorado 2016). Under the most realistic model of an *hs* relationship, we found that strongly deleterious mutations do accumulate at much higher frequencies in larger populations, leading to much faster extinction following contraction (Figure 4). However, we also found that simulations with an *hs* relationship in SLiM were extremely computationally intensive, such that we were unable to obtain results for the larger ancestral carrying capacities of 10,000 and 15,000 (see Supplementary Methods). Nevertheless, these results demonstrate a strong concordance between the *hs* relationship and our *hmix* model, suggesting that our *hmix* model captures the key features of the *hs* relationship (Figures 4 and S2).

**Figure 4:**
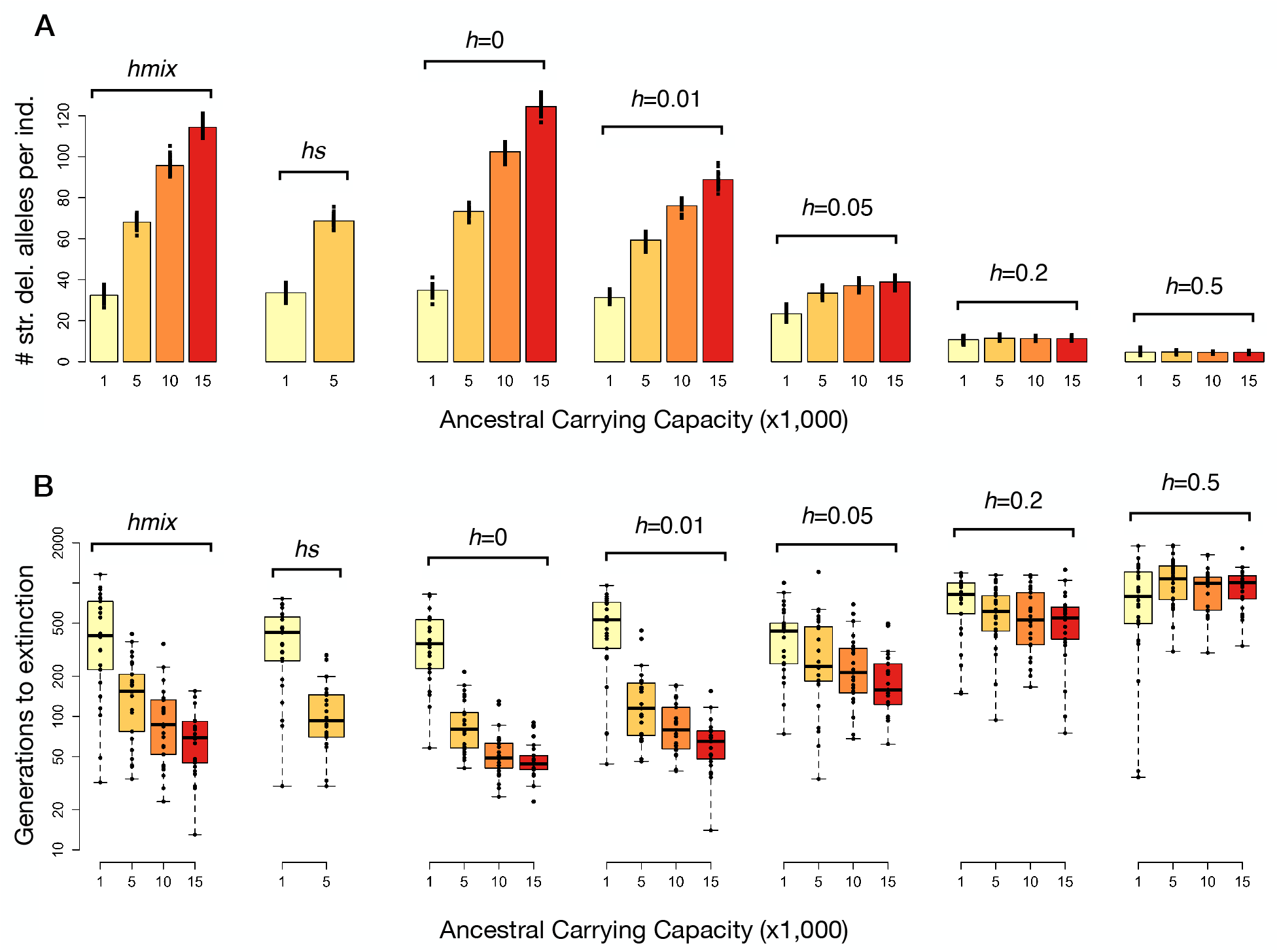
Impact of dominance coefficients on the accumulation of strongly deleterious mutations and extinction times following a contraction. (A) Mean number of strongly deleterious alleles per individual (*s* < −0.01) prior to contraction in ancestral populations of varying size and under varying models of dominance. (B) Time to extinction following contraction to K_endangered_ = 25 plotted on a log scale. Note that we were unable to obtain results for K_ancestral_ = 10,000 and 15,000 under the *hs* relationship due to computational limitations. For each parameter setting, 25 simulation replicates were run.

When assuming a single dominance coefficient for all mutations, our results demonstrate a range of outcomes that depended on the assumed dominance coefficient. Specifically, when assuming *h*=0 or 0.01, we again find that larger populations harbor higher levels of strongly deleterious variation, with strikingly similar patterns to the *hs* relationship or *hmix* model (Figure 4). However, this effect is greatly diminished when mutations are only partially recessive (*h*=0.05 or 0.2), and is nonexistent when mutations are additive (*h*=0.5), as expected (Figure 4). Notably, although the average dominance coefficient under our assumed *hs* relationship is approximately 0.2 (Agrawal and Whitlock 2011), our results with *h*=0.2 are dramatically different from those under the *hs* relationship (Figure 4).

To further investigate the importance of strongly vs weakly/moderately deleterious mutations in our *hmix* model, we ran simulations where we truncated our DFE to only permit strongly deleterious (*s* < −0.01 and *h*=0) or weakly/moderately deleterious mutations (*s* ≥ −0.01 and *h*=0.25) to enter the population. More specifically, in the case of permitting only strongly deleterious mutations, we allowed any mutation with *s* < −0.01 to enter the population as normal, however mutations with *s* ≥ −0.01 instead became neutral mutations. Here, our results for simulations that only included strongly deleterious mutations were notably similar to the full *hmix* model, with extinction times that depended strongly on ancestral demography (Figure S8). By contrast, results for simulations that only included weakly/moderately deleterious mutations were dramatically different from the full model, and had greatly increased extinction times (Figure S8). Given that strongly deleterious mutations constitute only ~25% of all new deleterious mutations under our assumed DFE (Kim et al. 2017), these results strikingly demonstrate their disproportionate impact on extinction risk relative to the effects of more weakly or moderately deleterious mutations.

As a final way of exploring the impact of recessive deleterious mutations, we also conducted simulations where we decreased the target size for deleterious mutations assuming *h*=0. The motivation for these simulations was to test whether we would still observe an impact of ancestral demography on extinction times when lowering the number of genes that could accumulate fully recessive deleterious mutations. To do this, we decreased the number of genes (*g*) in our simulations from 20,000 to *g* ∈ {1,000, 5,000, 10,000, and 15,000}. Here, we observed that the effect of ancestral demography is still present, though greatly diminished, with as few as 1,000 genes, and remains substantial with as few as 5,000 genes (Figure S9).

### Genetic rescue simulations

We next explored how demography and strongly deleterious variation impact the effectiveness of genetic rescue assuming an *hmix* model of dominance. Here, we quantify the effectiveness of genetic rescue as the extent to which the introduction of migrants to the endangered population increased extinction times. When translocating five migrants from one of four source populations (Figure 5A-B) to an endangered population with K=25, we found that all source populations led to increases in time to extinction relative to a no-rescue scenario (Figure 5C). Importantly, however, we found that the magnitude of increase in time to extinction was highly dependent on source population demography and levels of strongly deleterious variation. For example, whereas genetic rescue from the large source population (K=10,000) led to a notable increase in mean time to extinction of 16%, rescue from the moderate-sized source population resulted in a much more dramatic increase of 130% (Figure 5C). Genetic rescue from small and moderately-inbred populations (Figure S10) also resulted in increases in mean time to extinction that were at least as great as that of the large source population (63% increase for K=100, 13% increase for K=25) (Figure 5C). We observed similar patterns when conducting simulations with a larger endangered population (K=50) (Figure S11), though the beneficial effects of genetic rescue were somewhat diminished, likely due to the larger recipient population size when translocation was initiated (N ≤ 15) as well as the greater efficacy of purging within the larger recipient population.

**Figure 5:**
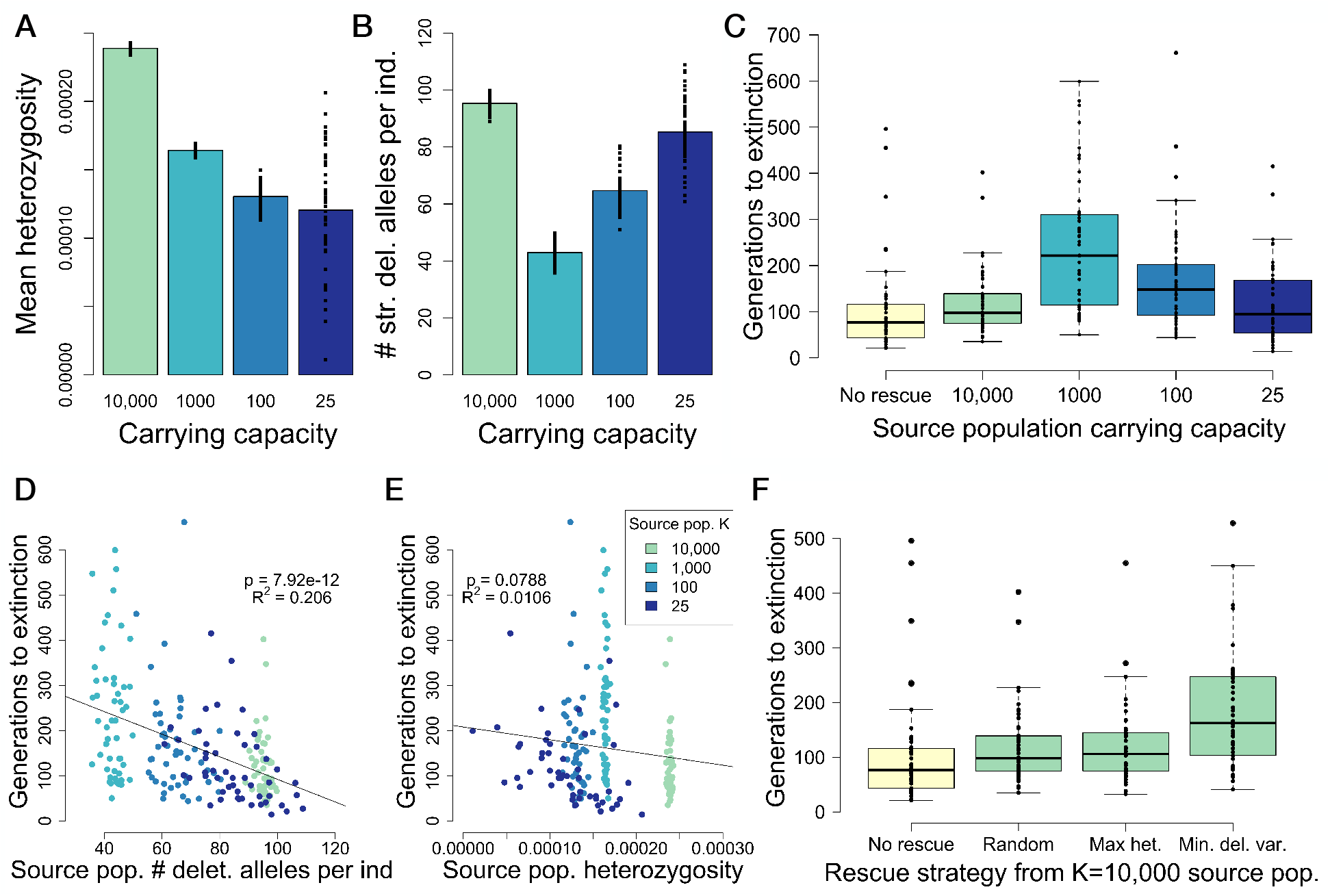
Source population deleterious variation determines the effectiveness of genetic rescue. (A) Average heterozygosity of source populations used for genetic rescue during the generation of rescue. (B) Mean number of strongly deleterious alleles (*s* < −0.01) per individual in the source populations used for genetic rescue. See Figure S10 for source population fitness and levels of inbreeding. (C) Time to extinction following genetic rescue from source populations of varying size. (D) Time to extinction following genetic rescue is negatively correlated with the number of recessive strongly deleterious alleles (*s* < −0.01) per individual used for rescue. (E) Time to extinction following genetic rescue is not correlated with the heterozygosity of the source population. See Figure S13 for regression results when considering only the K=25 source population. (F) Time to extinction under different strategies of genetic rescue from the K=10,000 source population. For each parameter setting, 50 simulation replicates were run.

Examination of individual replicates again offers insight into the factors driving extinction in these populations (Figures 6 and S12). In all simulated scenarios, we observed a large increase in fitness immediately following the introduction of migrants (Figures 6 and S12), likely due to the high strength of heterosis (masking of fixed recessive deleterious alleles) following initial admixture. In most cases, these increases in fitness led to population growth, though the extent to which population sizes increased was strongly influenced by environmental stochasticity (Figures 6 and S12). Soon after genetic rescue, however, the resumption of inbreeding again led to a decline in fitness (Figures 6 and S12). Here, the rate of fitness decline was determined by the levels of strongly deleterious variation in the migrant individuals and their descendants. For example, when translocation occurred from a large source population (K=10,000), levels of strongly deleterious variation remained high after genetic rescue, eventually resulting in severe inbreeding depression once inbreeding resumed (Figure 6). By contrast, when the moderate-sized source population was used (K=1,000), levels of strongly deleterious variation dramatically decreased following genetic rescue, greatly decreasing the severity of inbreeding depression in future generations (Figure 6).

**Figure 6:**
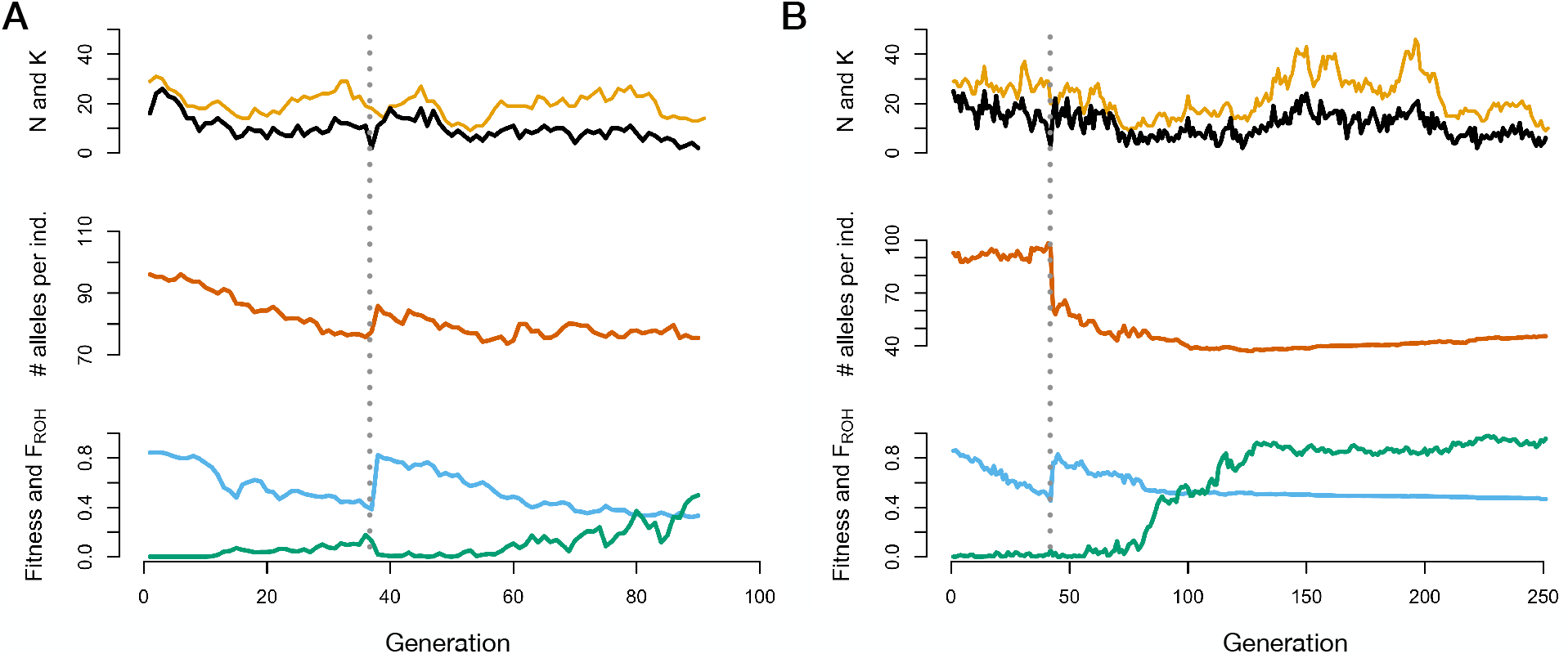
Representative examples of genetic rescue simulations with source populations of varying size. (A) Example when a large source population (K=10,000) is used for genetic rescue. (B) Example when a moderate-sized (K=1,000) is used. The top of each panel shows population size (N) in black and carrying capacity (K) in gold, the middle shows the average number of strongly deleterious alleles (*s* < −0.01) per individual, and the bottom shows mean absolute fitness in blue and mean inbreeding coefficient (F_ROH_) in green. The dashed grey line indicates the generation in which migration occurred. For both replicates, K_endangered_ = 25 and an *hmix* model of dominance was assumed. Note the differing y-axis scales for the middle panels and differing x-axis scales for panels A and B.

These results demonstrate that, although the larger source populations have higher fitness and genetic diversity, they also harbor a high number of heterozygous recessive strongly deleterious mutations (Figures 5 and S10). These mutations are quickly made homozygous by inbreeding in the endangered population following translocation, exacerbating the severity of inbreeding depression and eventually contributing to extinction. In support of this interpretation, we found that time to extinction following genetic rescue is predicted by the average number of strongly deleterious alleles per individual in the source population when examining all source populations simultaneously (Figure 5D). By contrast, we found that average source population heterozygosity was not very predictive of time to extinction, where in fact we observed a slight negative correlation due to the fact that source populations with higher genetic diversity also tend to harbor higher levels of recessive strongly deleterious variation (Figure 5E). We obtained similar results when restricting this analysis to the K=25 source population, the source population for which there was the highest variability in strongly deleterious variation and genetic diversity (Figure S13). However, in this case, we do see strong negative correlation between genetic diversity and extinction times, which is likely driven by heterozygosity being correlated with strongly deleterious variation at this intra-population scale.

Our finding that source population strongly deleterious variation predicts the efficacy of genetic rescue suggests that genome-wide levels of deleterious variation could be used to select individuals for genetic rescue. We explored the efficacy of this strategy by selecting the individuals with the fewest strongly deleterious alleles (*s* < −0.01) from the large source population (K=10,000) for rescue. This approach resulted in an increase in mean time to extinction of 74% compared to the non-rescue scenario, a 49% increase relative to randomly selecting individuals from the large source population (Figure 5F). By contrast, when individuals with the highest genome-wide heterozygosity were selected, we observed an increase in time to extinction that was no greater than when individuals were selected at random (Figure 5F). These results further support the causal relationship between strongly deleterious variation and extinction risk due to inbreeding depression, as well as the lack of this relationship between heterozygosity and extinction risk.

Lastly, we explored the effects of varying the number of migrants (1, 5, or 10) as well as the number of translocation events (1, 2, or 5), focusing on the K=10,000 and K=1,000 source populations. These simulations show persistent increases in time to extinction with increasing number of translocations regardless of the source population (Figure S14), suggesting that the efficacy of genetic rescue does not diminish with each additional migration event. When varying the number of migrants, we found that extinction times increased slightly in the case where migrants were drawn from the K=1,000 source population, but did not increase when drawing migrants from the K=10,000 source population (Figure S14).

## Discussion

Our simulations demonstrate the central importance of demographic history and recessive strongly deleterious mutations in determining the risk of extinction due to inbreeding depression in small and isolated populations. We find that populations that were historically large have a much higher risk of extinction following a population contraction compared to historically-smaller populations (Figure 2). These findings may be counterintuitive given the thinking that small populations should be less fit due to an accumulation of weakly deleterious alleles (Kimura et al. 1963; Lynch and Gabriel 1990; Lynch et al. 1995; Battaillon and Kirkpatrick 2000). The key insight that our simulations highlight is that larger ancestral populations harbor more recessive strongly deleterious mutations (Figure 2), due to these mutations being hidden from purifying selection as heterozygotes. The exposure of these mutations as homozygous by inbreeding in small populations can lead to dramatic reductions in fitness and drive rapid extinction, well before ‘mutational meltdown’ due to an accumulation of weakly deleterious mutations can occur (Lynch and Gabriel 1990; Lynch et al. 1995). By demonstrating that this effect is sufficient to ultimately drive extinction on short timescales, our simulations illustrate the importance of recessive deleterious variation as a mechanism of inbreeding depression (Charlesworth and Willis 2009; Hedrick and Garcia-Dorado 2016). Notably, we do not include any forms of heterozygote advantage in our model, which represents an alternative mechanism for inbreeding depression, though one that has received far less empirical support (Charlesworth and Willis 2009; Hedrick and Garcia-Dorado 2016). Thus, our finding that recessive deleterious mutations alone can drive rapid extinction in small populations serves as further support for these mutations being the primary cause of inbreeding depression.

Although the insight that strongly deleterious mutations that are fully recessive can accumulate at greater frequencies in larger populations has been noted elsewhere (Nei 1968; Hedrick 2002; Hedrick and Garcia-Dorado 2016; Szpiech et al. 2019), our results add to this work by demonstrating that that these effects persist under realistic genomic parameters and models of dominance, including both an *hs* relationship and our *hmix* model (Figure 4). This result is due to the fact that strongly deleterious mutations under these models tend to be highly recessive (Simmons and Crow 1977; Agrawal and Whitlock 2011; Huber et al. 2018), and are therefore hidden from purifying selection when present as heterozygotes in large populations Moreover, when examining models with a single dominance coefficient for all mutations, we find that our results with fully recessive mutations (*h*=0 or 0.01) are highly similar to those under more realistic models of dominance, whereas the impact of ancestral demography is greatly diminished with partially recessive (*h*=0.05 or 0.2) mutations and nonexistent with additive (*h*=0.5) mutations (Figure 4). Given that the mean dominance coefficient under our assumed *hs* relationship is 0.2 (Agrawal and Whitlock 2011), the strong dissimilarity between our results under an *hs* relationship and when assuming *h*=0.2 for all mutations is notable, and suggests that the key factor mediating extinction dynamics is the presence of fully recessive strongly deleterious mutations, rather than the overall mean dominance coefficient. In support of this interpretation, we show that simulations that only include the fraction of mutations that are strongly deleterious and fully recessive (*h*=0) exhibit extinction times that are highly similar to extinction times under an *hs* relationship or *hmix* model, whereas simulations that only include the fraction of mutations that are weakly or moderately deleterious and partially recessive (*h*=0.25) have much longer extinction times that depend only minimally on ancestral demography (Figure S8). This result is especially striking when considering that strongly deleterious mutations comprise only ~25% of the overall number of deleterious mutations under our assumed DFE (Kim et al. 2017). Taken together, these results further bolster our conclusion that strongly deleterious mutations are the primary determinant of extinction risk due to inbreeding depression in small populations. However, gaining a more precise understanding of these dynamics will greatly depend on improving our knowledge of the distribution of dominance coefficients across various taxa.

The considerable influence of ancestral demography on extinction risk that our simulations reveal has important implications for assessing the threat of extinction due to inbreeding depression in wild populations. Quantifying inbreeding depression in the wild and predicting the threat it poses to extinction remains a major challenge in conservation biology, and the reasons why some small populations suffer from inbreeding depression and others do not is often unclear (Hedrick and Garcia-Dorado 2016). Our simulations demonstrate that these differences may be determined in large part by the ancestral demography of a species. Consequently, we suggest that information on ancestral demography, which is increasingly becoming accessible using genomic data (Beichman et al. 2018), should be more widely incorporated into extinction risk predictions. In particular, given that nearly all threatened populations have recently undergone reductions in size due to anthropogenic pressures, these results suggest that their continued persistence may depend crucially on their ancestral demography and load of recessive strongly deleterious mutations. In addition, our results also suggest that island populations that have historically been small may in fact have reduced extinction risk due to genetic factors due to their high isolation and smaller population size, which may have facilitated purging of recessive strongly deleterious mutations (Laws and Jamieson 2011; Robinson et al. 2018). However, our simulations also reveal that the fate of small populations is highly stochastic, and that even under the same ecological and genetic parameters, time to extinction can vary substantially (Figures 2 and 4). Thus, predictions of extinction risk for any wild population should necessarily be accompanied by a fair amount of uncertainty.

Our results also provide insight on how best to conduct a genetic rescue, which is becoming increasingly necessary for maintaining small and isolated populations under growing anthropogenic pressures (Whiteley et al. 2015; Hedrick and Garcia-Dorado 2016; Frankham et al. 2017; Bell et al. 2019). Consistent with many other studies (e.g., Hogg et al. 2006; Johnson et al. 2010; Frankham 2015; Åkesson et al. 2016), our simulations highlight the beneficial effects of genetic rescue, supporting recent calls for its more widespread application (Whiteley et al. 2015; Frankham et al. 2017; Bell et al. 2019). However, in stark contrast to existing recommendations (Pickup et al. 2012; Whiteley et al. 2015), we found that targeting large source populations with high genetic diversity was among the least effective genetic rescue strategies (Figures 5 and S11). Instead, our results demonstrate that the effectiveness of genetic rescue can potentially be maximized by drawing migrants from moderate-sized source populations which have fewer strongly deleterious mutations (Figures 5 and S11). These source populations are ideal due to being small enough to carry few strongly deleterious recessive mutations in the heterozygous state, but not so small that they accumulate a substantial load of fixed weakly or moderately deleterious mutations. Finally, our results demonstrate that even small and somewhat inbred populations can also serve as effective source populations (Figures 5C and S10), as other studies have shown (Heber et al. 2012b,a).

Although our results suggest that large populations may not be ideal source populations for genetic rescue, we demonstrate that the effectiveness of this strategy can be greatly increased when individuals are screened for low levels of strongly deleterious variation (Figure 5F). However, applications of such approach may remain limited by our ability to accurately predict the fitness consequences of individual mutations, which remains a major challenge even in humans (Eilbeck et al. 2017). Our results also demonstrate that repeated genetic rescue from large or moderate-sized source populations can result in persistent beneficial effects (Figure S14), highlighting the efficacy of this strategy for populations that are destined to remain small and isolated. One inevitable consequence of this or any other genetic rescue strategy, however, is a loss of native ancestry (Johnson et al. 2010; Adams et al. 2011; Harris et al. 2019), which can potentially swamp out locally adapted alleles (though see Fitzpatrick et al. 2020). Although we do not track levels of admixture in our simulations, it is probable that the post-rescue populations are composed of highly admixed individuals (Harris et al. 2019).

In addition to providing guidelines for how to best conduct a genetic rescue, our results also have implications for understanding the mechanisms underlying genetic rescue. Two mechanisms have generally been proposed for genetic rescue: heterosis (masking of fixed recessive deleterious mutations) and adaptive evolution (increases in fitness due to selection on new alleles) (Whiteley et al. 2015). By demonstrating that, even in the absence of adaptive variation, migration into small and inbred populations can lead to increases in fitness and demographic rates consistent with those observed in empirical systems (e.g., Hogg et al. 2006; Johnson et al. 2010; Adams et al. 2011; Åkesson et al. 2016), our results suggest that heterosis alone may be able to explain much of the beneficial effects of genetic rescue. However, a more thorough investigation of the relative roles of heterosis and adaptive evolution as mechanisms of genetic rescue could be conducted by incorporating adaptive mutations into our simulation framework. Finally, our finding that increases in fitness following migration do not always lead to increases in population size due to the effects of environmental stochasticity (Figure S12) suggests that definitions of genetic rescue based on demographic effects alone may be restrictively narrow, as other authors have suggested (Hedrick et al. 2011). In particular, this result has relevance to the debate of whether the Isle Royale wolf population truly experienced a genetic rescue, given that the population size did not increase substantially following migration, perhaps due to poor environmental conditions (Adams et al. 2011; Hedrick et al. 2011)

Our simulation framework has several notable limitations. First, due to the high computational load of our simulations (see Supplementary Methods), we were unable to examine ancestral populations with carrying capacities larger than 15,000, limiting the observed impact of ancestral demography. Second, we did not examine heterozygote advantage as a potential mechanism for inbreeding depression, which could impact the negative relationship between extinction times and source population heterozygosity that we observed (Figure 5E). However, empirical support for heterozygote advantage as a mechanism for inbreeding depression is scarce (Charlesworth and Willis 2009; Hedrick and Garcia-Dorado 2016), suggesting that it may not substantially impact our results. In addition, we did not include adaptive mutations in our model, which may also be relevant to the ability of small populations to persist under environmental change, as well as for exploring the impacts of outbreeding depression due to differential local adaptation. Although recent work has suggested that concerns about outbreeding depression may have been overstated (Whiteley et al. 2015; Frankham et al. 2017), assessing its influence in determining source population selection nevertheless represents an important area for future research. Finally, we emphasize that the demographic scenarios explored here may not be applicable to all small and isolated populations. Specifically, our assumption of an instantaneous contraction from a large ancestral population to a small endangered population may not capture the demographic trajectory of many populations that have experienced more gradual declines. In these cases, a gradual decline may have facilitated purging of strongly deleterious mutations (Figure S5), decreasing the necessity for genetic rescue (e.g., Xue et al. 2015; Robinson et al. 2018). Overall, we recommend that any conservation actions motivated by our results carefully consider the demography of the population of interest, ideally using simulations.

In conclusion, our results highlight the detrimental effects of strongly deleterious variation in small populations, and suggest that many conservation strategies for endangered species may be improved by minimizing strongly deleterious variation rather than maximizing heterozygosity. These results are especially relevant to the ongoing reintroduction of wolves to Isle Royale, which has been guided by the goal of maximizing the genetic diversity of the new population. Although our simulations are motivated by this example, by examining a wide range of demographic scenarios, our results have implications beyond the Isle Royale wolf population. Future research should explore implementation of these strategies by expanding the use of genomic tools and assessments of deleterious variation in the context of the conserving small and isolated populations (Fitzpatrick and Funk 2019). Given the great expense of most translocation programs, incorporating genomic approaches represents a sound investment with the potential to substantially postpone the need for future intervention.

## Acknowledgements

We are grateful to Jacqueline Robinson, Bernard Kim, Brad Shaffer, Marty Kardos, and members of the Lohmueller and Wayne labs for helpful feedback and ideas. We thank Ben Haller for assistance with SLiM. This research was supported by NIH grant R35GM119856 (to K.E.L.).

## Author Contributions

CCK and KEL conceived the project, CCK ran the simulations and analyzed the results, all authors contributed to writing and editing the manuscript.

## Data Accessibility

Simulation and data processing scripts are available at https://github.com/ckyriazis/genetic_rescue_simulations.

## References

Adams, J. R., L. M. Vucetich, P. W. Hedrick, R. O. Peterson, and J. A. Vucetich. 2011. Genomic sweep and potential genetic rescue during limiting environmental conditions in an isolated wolf population. Proc. R. Soc. B Biol. Sci. 278:3336–3344.

Agrawal, A. F., and M. C. Whitlock. 2011. Inferences about the distribution of dominance drawn from yeast gene knockout data. Genetics 187:553–566.

Åkesson, M., O. Liberg, H. Sand, P. Wabakken, S. Bensch, and Ø. Flagstad. 2016. Genetic rescue in a severely inbred wolf population. Mol. Ecol. 25:4745–4756.

Allendorf, F. W., and R. F. Leary. 1986. Heterozygosity and fitness in natural populations of animals. Pp. 58–72 *in* Conservation biology: the science of scarcity and diversity.

Battaillon, T., and M. Kirkpatrick. 2000. Inbreeding depression due to mildly deleterious mutations in finite populations: size does matter. Genet. Res. (Camb). 75:75–81.

Beichman, A. C., E. Huerta-Sanchez, and K. E. Lohmueller. 2018. Using Genomic Data to Infer Historic Population Dynamics. Annu. Rev. Ecol. Evol. Syst. 49:433–456.

Bell, D. A., Z. L. Robinson, W. C. Funk, S. W. Fitzpatrick, F. W. Allendorf, D. A. Tallmon, and A. R. Whiteley. 2019. The Exciting Potential and Remaining Uncertainties of Genetic Rescue. Trends Ecol. Evol. 1–11. Elsevier Ltd.

Caballero, A., I. Bravo, and J. Wang. 2017. Inbreeding load and purging: Implications for the short-term survival and the conservation management of small populations. Heredity (Edinb). 118:177–185. Nature Publishing Group.

Caughley, G. 1994. Directions in Conservation Biology. J. Anim. Ecol. 63:215–244.

Charlesworth, D., and J. H. Willis. 2009. The genetics of inbreeding depression. Nat. Rev. Genet. 10:783–796.

Eilbeck, K., A. Quinlan, and M. Yandell. 2017. Settling the score: Variant prioritization and Mendelian disease. Nat. Rev. Genet. 18:599–612. Nature Publishing Group.

Fitzpatrick, S. W., G. S. Bradburd, C. T. Kremer, P. E. Salerno, L. M. Angeloni, and W. C. Funk. 2020. Genomic and Fitness Consequences of Genetic Rescue in Wild Populations. Curr. Biol. 30:1–6. Elsevier Ltd.

Fitzpatrick, S. W., and W. C. Funk. 2019. Genomics for Genetic Rescue. P. *in* Wildlife Conservation Genomics.

Frankham, R. 2015. Genetic rescue of small inbred populations: meta-analysis reveals large and consistent benefits of gene flow. Mol. Ecol. 24:2610–2618.

Frankham, R., J. D. Ballou, K. Ralls, M. Eldridge, M. R. Dudash, C. B. Fenster, R. C. Lacy, and P. Sunnucks. 2017. Genetic Management of Fragmented Animal and Plant Populations. Oxford University Press, Oxford, UK.

Grossen, C., F. Guillaume, L. F. Keller, and D. Croll. 2020. Purging of highly deleterious mutations through severe bottlenecks in Alpine ibex. Nat. Commun. 11:1001. Springer US.

Haller, B. C., and P. W. Messer. 2019. SLiM 3: Forward Genetic Simulations Beyond the Wright-Fisher Model. Mol. Biol. Evol. 36:632–637.

Harris, K., Y. Zhang, and R. Nielsen. 2019. Genetic rescue and the maintenance of native ancestry. Conserv. Genet. 20:59–64. Springer Netherlands.

Heber, S., J. V. Briskie, and L. A. Apiolaza. 2012a. A test of the “genetic rescue” technique using bottlenecked donor populations of Drosophila melanogaster. PLoS One 7.

Heber, S., A. Varsani, S. Kuhn, A. Girg, B. Kempenaers, and J. Briskie. 2012b. The genetic rescue of two bottlenecked South Island robin populations using translocations of inbred donors. Proc. R. Soc. B Biol. Sci. 280.

Hedrick, P. W. 2002. Lethals in Finite Populations. Evolution (N. Y). 56:654–657.

Hedrick, P. W., J. R. Adams, and J. A. Vucetich. 2011. Reevaluating and Broadening the Definition of Genetic Rescue. Conserv. Biol. 25:1069–1070.

Hedrick, P. W., and A. Garcia-Dorado. 2016. Understanding Inbreeding Depression, Purging, and Genetic Rescue. Trends Ecol. Evol. 31:940–952. Elsevier Ltd.

Hedrick, P. W., R. O. Peterson, L. M. Vucetich, J. R. Adams, and J. A. Vucetich. 2014. Genetic rescue in Isle Royale wolves: genetic analysis and the collapse of the population. Conserv. Genet. 15:1111–1121.

Hedrick, P. W., J. A. Robinson, R. O. Peterson, and J. A. Vucetich. 2019. Genetics and extinction and the example of Isle Royale wolves. Anim. Conserv. 22:302–309.

Henn, B. M., L. R. Botigué, S. Peischl, I. Dupanloup, M. Lipatov, B. K. Maples, A. R. Martin, S. Musharoff, H. Cann, M. P. Snyder, L. Excoffier, J. M. Kidd, and C. D. Bustamante. 2016. Distance from sub-Saharan Africa predicts mutational load in diverse human genomes. Proc. Natl. Acad. Sci. 113:E440–E449.

Hogg, J. T., S. H. Forbes, B. M. Steele, and G. Luikart. 2006. Genetic rescue of an insular population of large mammals. Proc. R. Soc. B 273:1491–1499.

Huber, C. D., A. Durvasula, and A. M. Hancock. 2018. Gene expression drives the evolution of dominance. Nat. Commun. 9:1–11. Springer US.

Huber, C. D., B. Y. Kim, C. D. Marsden, and K. E. Lohmueller. 2017. Determining the factors driving selective effects of new nonsynonymous mutations. Proc. Natl. Acad. Sci. 114:4465–4470.

Johnson, W. E., D. P. Onorato, M. E. Roelke, E. D. Land, M. Cunningham, R. C. Belden, R. McBride, D. Jansen, M. Lotz, D. Shindle, J. G. Howard, D. E. Wildt, L. M. Penfold, J. A. Hostetler, M. K. Oli, and S. J. O’Brien. 2010. Genetic restoration of the Florida panther. Science (80-.). 329:1641–1645.

Kim, B. Y., C. D. Huber, and K. E. Lohmueller. 2017. Inference of the Distribution of Selection Coefficients for New Nonsynonymous Mutations Using Large Samples. Genetics 206:345–361.

Kimura, M., T. Maruyama, and J. F. Crow. 1963. The mutation load in small populations. Genetics 1303–1312.

Lande, R. 1994. Risk of population extinction from fixation of new deleterious mutations. Evolution (N. Y). 48:1460–1469.

Laws, R. J., and I. G. Jamieson. 2011. Is lack of evidence of inbreeding depression in a threatened New Zealand robin indicative of reduced genetic load? Anim. Conserv. 14:47–55.

Lindblad-Toh, K., C. M. Wade, T. S. Mikkelsen, E. K. Karlsson, D. B. Jaffe, M. Kamal, M. Clamp, J. L. Chang, E. J. Kulbokas, M. C. Zody, E. Mauceli, X. Xie, M. Breen, R. K. Wayne, E. A. Ostrander, C. P. Ponting, F. Galibert, D. R. Smith, P. J. DeJong, E. Kirkness, P. Alvarez, T. Biagi, W. Brockman, J. Butler, C. W. Chin, A. Cook, J. Cuff, M. J. Daly, D. DeCaprio, S. Gnerre, M. Grabherr, M. Kellis, M. Kleber, C. Bardeleben, L. Goodstadt, A. Heger, C. Hitte, L. Kim, K. P. Koepfli, H. G. Parker, J. P. Pollinger, S. M. J. Searle, N. B. Sutter, R. Thomas, C. Webber, and E. S. Lander. 2005. Genome sequence, comparative analysis and haplotype structure of the domestic dog. Nature 438:803–819.

Lynch, M., I. J. Conery, and R. Burger. 1995. Mutation accumulation and the extinction of small populations. Am. Nat. 146:489–518.

Lynch, M., and W. Gabriel. 1990. Mutation load and the survival of small populations. Evolution (N. Y). 44:1725–1737.

McLaren, B. E., and R. O. Peterson. 1994. Wolves, Moose, and Tree Rings on Isle Royale. Science (80-.). 266:1555–1558.

Mech, L. D. 1966. The Wolves of Isle Royale.

Nei, M. 1968. The frequency distribution of lethal chromosomes in finite populations. Proc. Natl. Acad. Sci. 60:517–524.

O’Grady, J. J., B. W. Brook, D. H. Reed, J. D. Ballou, D. W. Tonkyn, and R. Frankham. 2006. Realistic levels of inbreeding depression strongly affect extinction risk in wild populations. Biol. Conserv. 133:42–51.

Peterson, R. O., R. E. Page, and K. M. Dodge. 1984. Wolves, Moose, and the Allometry of Population Cycles. Science (80-.). 224:1350–1352.

Pickup, M., D. L. Field, D. M. Rowell, and A. G. Young. 2012. Source population characteristics affect heterosis following genetic rescue of fragmented plant populations. Proc. R. Soc. B 280:1–9.

Reed, D. H., and R. Frankham. 2003. Correlation between Fitness and Genetic Diversity. Conserv. Biol. 17:230–237.

Robinson, J. A., C. Brown, B. Y. Kim, K. E. Lohmueller, and R. K. Wayne. 2018. Purging of Strongly Deleterious Mutations Explains Long-Term Persistence and Absence of Inbreeding Depression in Island Foxes. Curr. Biol. 28:1–8. Elsevier Ltd.

Robinson, J. A., D. Ortega-Del Vecchyo, Z. Fan, B. Y. Kim, B. M. Vonholdt, C. D. Marsden, K. E. Lohmueller, and R. K. Wayne. 2016. Genomic Flatlining in the Endangered Island Fox. Curr. Biol. 26:1183–1189. Elsevier Ltd.

Robinson, J. A., J. Räikkönen, L. M. Vucetich, J. A. Vucetich, R. O. Peterson, K. E. Lohmueller, and R. K. Wayne. 2019. Genomic signatures of extensive inbreeding in Isle Royale wolves, a population on the threshold of extinction. Sci. Adv. 5:1–13.

Simmons, M. J., and J. F. Crow. 1977. Mutations affecting fitness in Drosophila populations. Ann. Rev. Genet. 11:49–78.

Spielman, D., B. W. Brook, and R. Frankham. 2004. Most species are not driven to extinction before genetic factors impact them. Proc. Natl. Acad. Sci. 101:15261–15264.

Szpiech, Z. A., A. C. Mak, M. J. White, D. Hu, C. Eng, E. G. Burchard, and R. D. Hernandez. 2019. Ancestry-dependent Enrichment of Deleterious Homozygotes in Runs of Homozygosity. Am. J. Hum. Genet. 105:1–16. ElsevierCompany.

Van Der Valk, T., M. De Manuel, T. Marquez-Bonet, and K. Guschanski. 2019. Estimates of genetic load suggest extensive genetic purging in mammalian populations. bioRxiv.

Wayne, R., N. Lehman, D. Girman, P. Gogan, D. Gilbert, K. Hansen, R. Peterson, U. Seal, A. Eisenhawer, L. Mech, and R. J. Krumenaker. 1991. Conservation Genetics of the Endangered Isle Royale Gray Wolf. Conserv. Biol. 5:41–51.

Whiteley, A. R., S. W. Fitzpatrick, W. C. Funk, and D. A. Tallmon. 2015. Genetic rescue to the rescue. Trends Ecol. Evol. 30:42–49. Elsevier Ltd.

Xue, Y., J. Prado-Martinez, P. H. Sudmant, V. Narasimhan, Q. Ayub, M. Szpak, P. Frandsen, Y. Chen, B. Yngvadottir, D. N. Cooper, M. De Manuel, J. Hernandez-rodriguez, I. Lobon, H. R. Siegismund, L. Pagani, M. A. Quail, C. Hvilsom, A. Mudakikwa, E. E. Eichler, M. R. Cranfield, and T. Marques-bonet. 2015. Mountain gorilla genomes reveal the impact of long-term population decline and inbreeding. Science (80-.). 348:242–245.

