## Supplemental Text and Figures for "Strongly deleterious mutations are a primary determinant of extinction risk due to inbreeding depression"

### Supplementary Methods

#### 1. Relating K to $N_e$ in the SLiM nonWF model

Translating the carrying capacity of a SLiM nonWF simulation to the corresponding Wright-Fisher effective population size is not entirely straightforward, for three reasons. First, when deleterious mutations are simulated, the size of the simulated population decreases relative to its carrying capacity, reflecting the decline in fitness of the population due to its mutational load (Table S1). Second, the stochastic fluctuations in the population size lead to a further reduction in its effective size relative to the census size. Finally, the use of overlapping generations in this model also contributes to a reduction in effective population size relative to census size. Overall, we find that the effective size of a population given our simulation parameters is approximately 70% of its carrying capacity, as estimated from the neutral heterozygosity of these populations using the expectation under the Wright-Fisher model that $N_e = \pi/4\mu$ (Table S1).

**Table S1** Carrying capacity, population size, and effective population size

| Carrying capacity | N at equilibrium with deleterious mutations* | Neutral heterozygosity ($\pi$)** | Ne ($\pi/4\mu$) | Ne/K |
|---|---|---|---|---|
| 15,000 | 13,367 | 4.22E-04 | 10551 | 0.703 |
| 10,000 | 8,891 | 2.80E-04 | 6988.88 | 0.699 |
| 5,000 | 4,394 | 1.41E-04 | 3524.23 | 0.705 |
| 1,000 | 876 | 2.82E-05 | 704.689 | 0.705 |

*****Averaged from 5 replicates with deleterious variation
**Averaged from 5 replicates without deleterious variation

**2. Additional details on our assumed distribution of fitness effects**

The distribution of fitness effects (DFE) we assumed in our simulations was inferred by Kim et al. 2017 using human polymorphism data. Depending on the specific dataset and assumed functional form of the DFE, they estimate that 24.5–29.8% of new nonsynonymous mutations are strongly deleterious. This estimate is somewhat lower than other similar studies (e.g., Boyko et al. 2008 estimated this figure to be 35.5%), though is likely to be a better estimate due to the much larger sample sizes used in Kim et al. 2017, which translates to greater power in detecting rare recessive strongly deleterious mutations.

The DFE inferred by Kim et al. 2017 assumed that all mutations are additive ($h$=0.5). Although our simulations generally assume that mutations are partially or fully recessive, we are not aware of any DFEs that have been inferred for a mammalian species under a (partially) recessive model. A DFE including partially recessive mutations would likely have >25% of strongly deleterious recessive mutations because the strength of selection needs to be greater to have the same impact on polymorphism when deleterious mutations are recessive compared to when they are additive. Thus, it is likely that our simulations are in fact conservative with regard to the number of strongly deleterious mutations segregating in natural populations. We suggest that future work should explore this issue in greater detail, though the qualitative results of our simulations are likely to remain the same.

**3. Discussion of computational limitations**

Forward-in-time population genetic simulations are often quite computationally intensive, and we found this to be the case for many of the simulations we ran in this study. For example, simulations with $K_{ancestral}$=10,000 typically took ~4 days and up to 48G of memory to complete, and simulations with $K_{ancestral}$=15,000 typically took nearly two weeks and up to 96G of memory to complete. The bulk of this computational time is during the burn-in, which grows rapidly with increasing $K_{ancestral}$ due both to the increasing number of generations for the burn-in (10*$K_{ancestral}$) as well as the larger number of individuals and mutations in larger populations. These computational times were further increased when we implemented an $hs$ relationship model of dominance, due to need to manually set the fitness effects of heterozygous genotypes every generation. For example, we found that $K_{ancestral}$=10,000 simulations that would take ~4 days to complete with a single dominance coefficient for all mutations instead took >2 weeks to complete, making them computationally intractable.

To speed up forward-in-time simulations, several approaches are typically employed, none of which were effective or appropriate in our situation. One common approach consists of rescaling the number of individuals simulated to be smaller than population size, and then adjusting other simulation parameters (mutation rates, recombination rates, and selection coefficients) accordingly, conserving the same values of the scaled parameters. Unfortunately, this approach is not appropriate in our case due to the need to model the true (i.e., unscaled) census size of a population for the purposes of the ecological dynamics in our model. Another approach that has recently become available in SLiM is the use of tree sequence recording, in which neutral mutations are initially omitted and later overlaid on a genealogy that is recorded for the simulation (Haller et al. 2019). However, this approach only becomes a faster alternative when long chromosomal segments with many neutral mutations are simulated (Haller et al. 2019), which is not the case for our simulations. One final option for speeding up simulations in SLiM is to remove fixed mutations, which may be appropriate when only the relative (rather than absolute) fitness of the simulated populations is of interest. Given our need to track absolute fitness for the purpose of modeling extinction due to inbreeding depression, this approach was also not appropriate in our case.

**4. Implementing the *hmix* model of dominance in SLiM**

To implement the *hmix* dominance model approach in SLiM (Figure S4), we created separate weakly/moderately and strongly deleterious mutation types, both of which draw new mutations from the same underlying DFE. Here, however, we ‘rejected’ mutations outside of the specified selection coefficient range by changing them to neutral. For example, a mutation drawn from the ‘strongly deleterious’ mutation type would enter the population if $s < -0.01$, but would be rejected if $s \geq -0.01$ by changing its selection coefficient to 0. Under this approach, half of all new deleterious mutations therefore become neutral mutations (~75% of strongly deleterious mutations and ~25% of weakly/moderately deleterious mutations), resulting in a ratio of deleterious to neutral mutations of 1:1 that is much lower than the desired ratio of 2.31:1. To account for this deficit of deleterious mutations, we increased the mutation rate by a factor of (2.31/3.31)/0.5=1.396. This also led to an increase in the number of neutral mutations in these simulations, however these mutations do not impact fitness and thus do not influence the underlying behavior of our model.

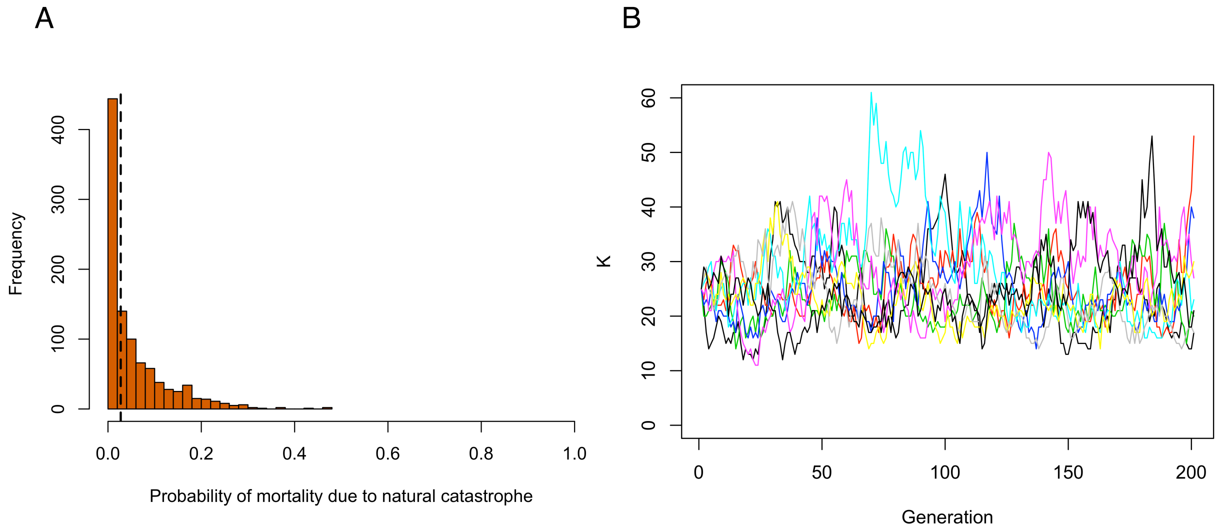


**Figure S1** Summary of stochastic population dynamics. (A) Histogram of 5000 simulations from a beta distribution with $\alpha = 0.5$ and $\beta = 8$, from which the probability of stochastic mortality due to a natural catastrophe in each generation of the simulation was drawn. The median is shown with a dashed line. (B) Ten simulated trajectories of stochastic Ornstein-Uhlenbeck model used to determine carrying capacity of the endangered population in simulations with $K_{mean} = 25$.

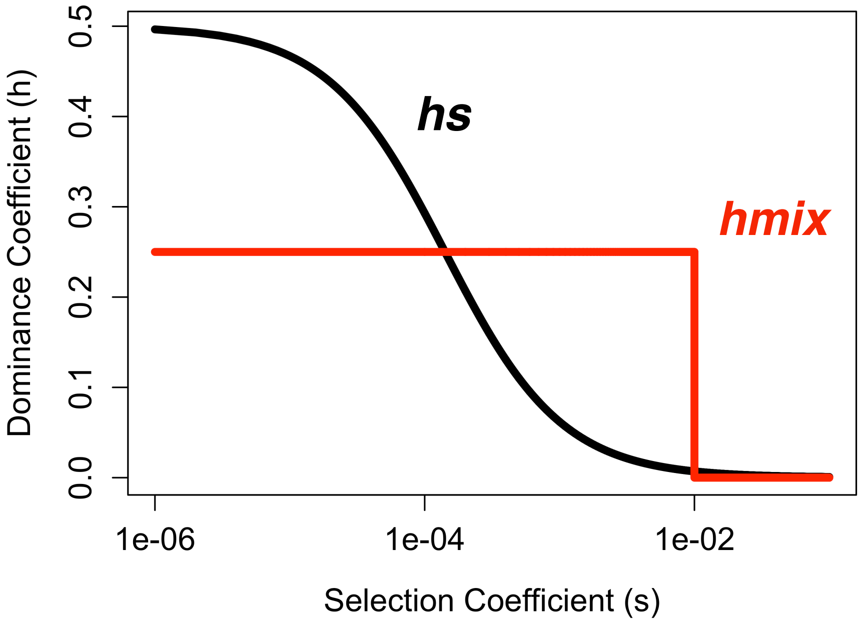


**Figure S2** Comparison of distribution of dominance coefficients under *hs* relationship from Agrawal and Whitlock (2011) and *hmix* model. Note that the x-axis is on a log scale.

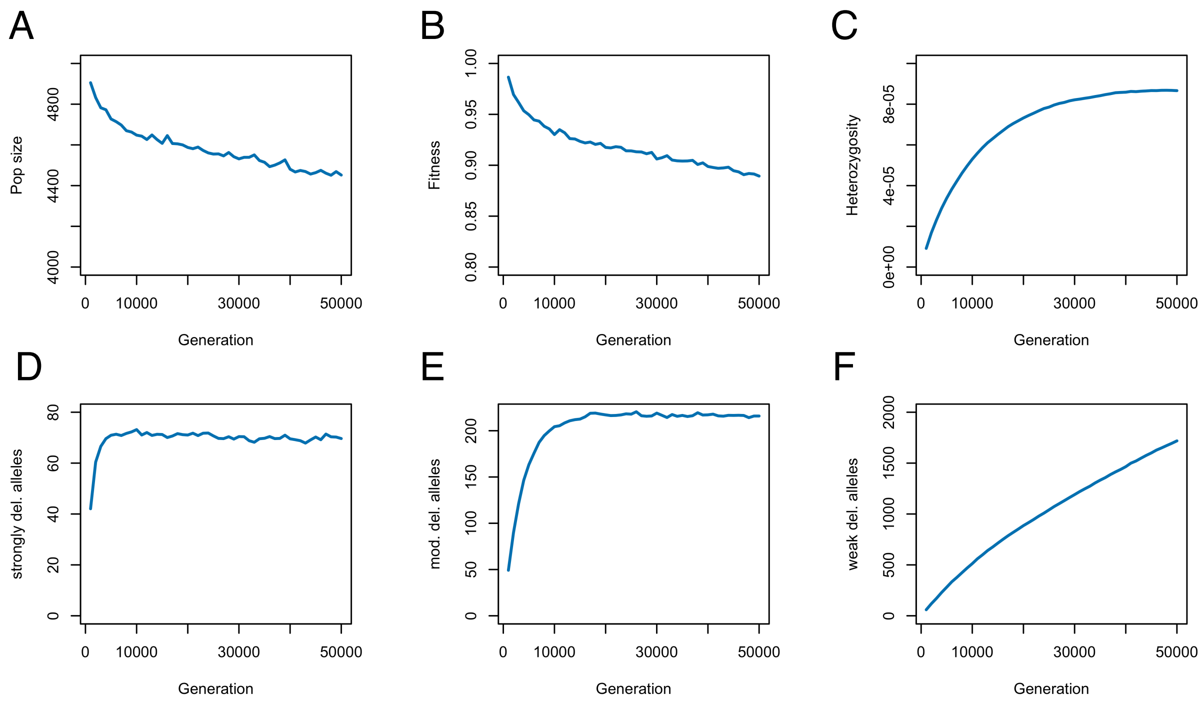


**Figure S3** Burn-in dynamics for the K=5000 population assuming $h$=0. (A) Population size, (B) mean absolute fitness, (C) mean heterozygosity, (D) average number of strongly deleterious ($s \leq$ -0.01) per individual, (E) average number of moderately deleterious (-0.01 $\leq s <$ -0.001) alleles per individual, (F) average number of weakly deleterious (-0.00001 $\leq s <$ -0.0001) alleles per individual. Statistics sampled every 1000 generations during the burn in.

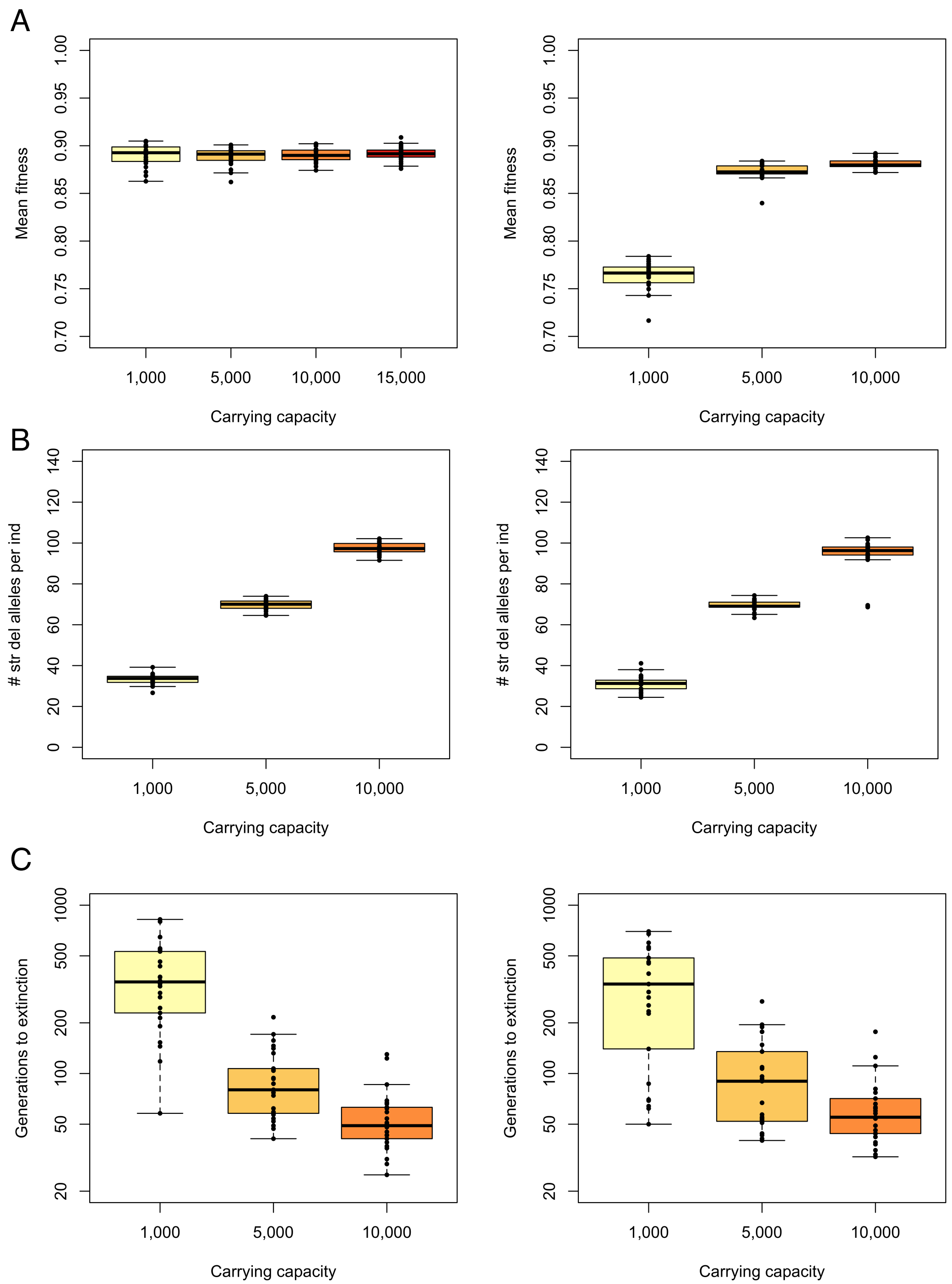


**Figure S4** Comparison of extinction dynamics between population contraction simulations with burn-in duration of 10*$K_{ancestral}$ (left panels) and simulations with burn-in duration of 10*$K_{ancestral}$ + 20,000 (right panels) assuming *h*=0. (A) Mean fitness prior to contraction. (B) Number of strongly deleterious alleles per individual ($s \leq -0.01$) prior to the contraction. (C) Time to extinction following contraction to $K_{endangered}$ = 25. Note that, although the addition of 20,000 generations to the burn in decreases fitness for the smaller populations, levels of strongly deleterious variation and extinction times remain unchanged.

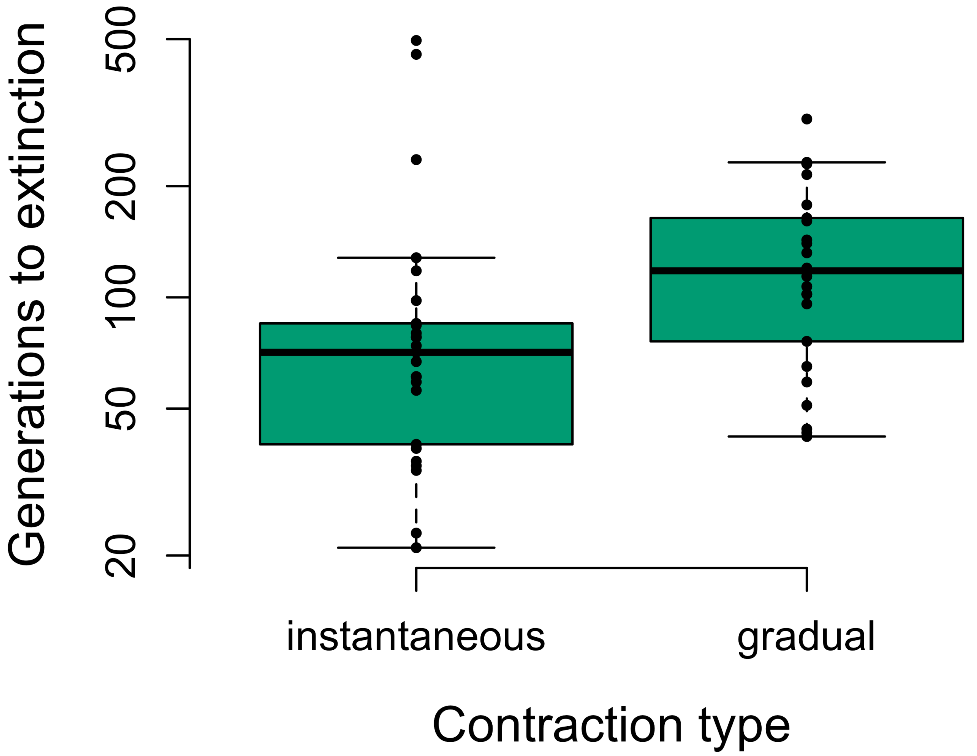


**Figure S5** Comparison of the time to extinction in an instantaneous vs gradual contraction from $K_{ancestral}$=10,000 to $K_{endangered}$=25 under an *hmix* model of dominance. For the gradual contraction scenario, the ancestral population first contracted to a carrying capacity of 1,000 for 200 generations before finally contracting to $K_{endangered}$=25. Note the longer extinction times for the gradual contraction, suggesting that it facilitated purging of strongly deleterious mutations. For each contraction type, 25 simulation replicates were run.

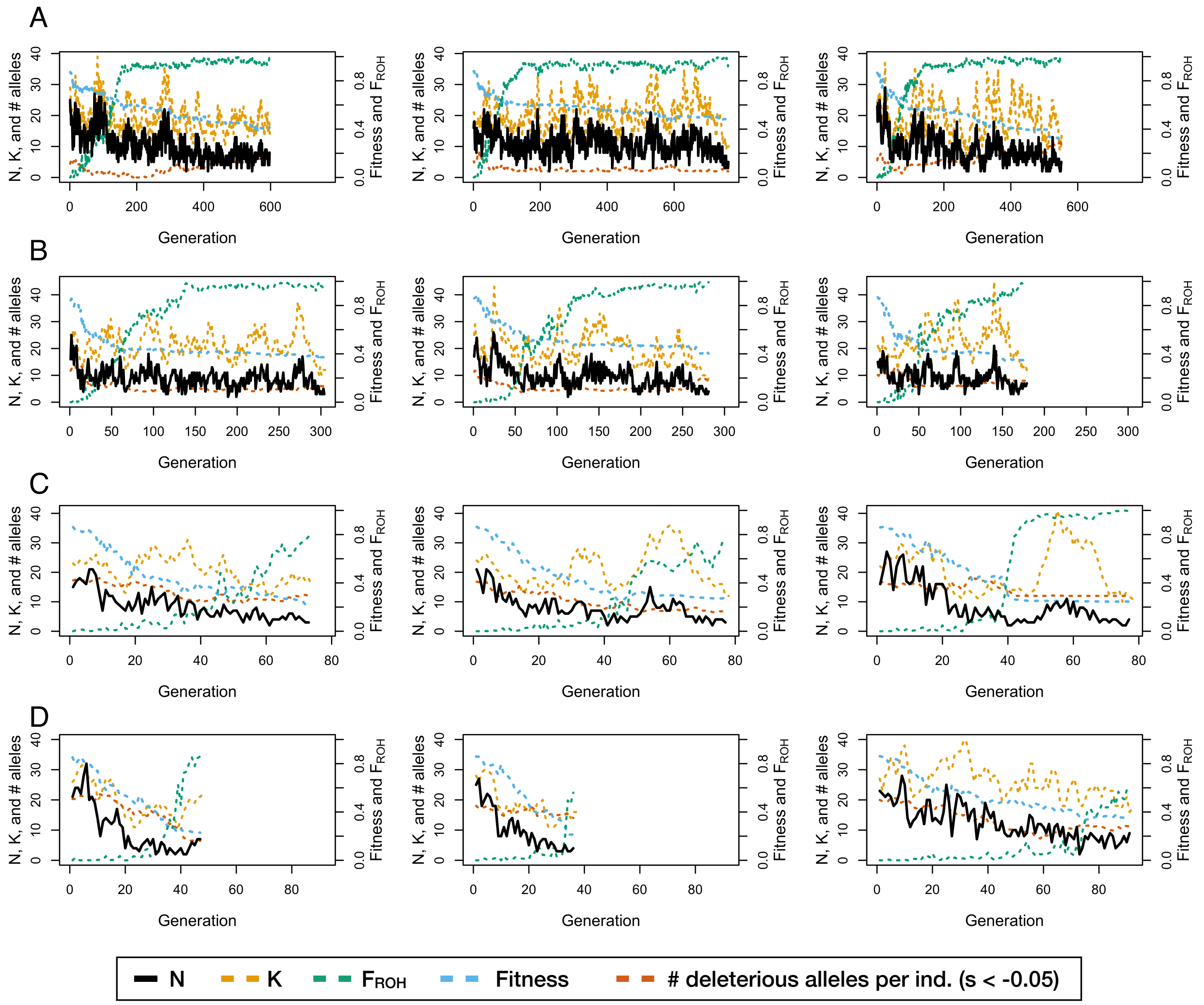


**Figure S6** Population trajectory of individual simulation replicates following contraction to endangered carrying capacity of 25 from ancestral carrying capacity of (A) 1,000, (B) 5,000, (C) 10,000, and (D) 15,000 under an *hmix* model of dominance. Three representative replicates are shown for each combination of parameters. Note the quicker time to extinction (denoted by the shorter x-axis) for the replicates with larger ancestral carrying capacities.

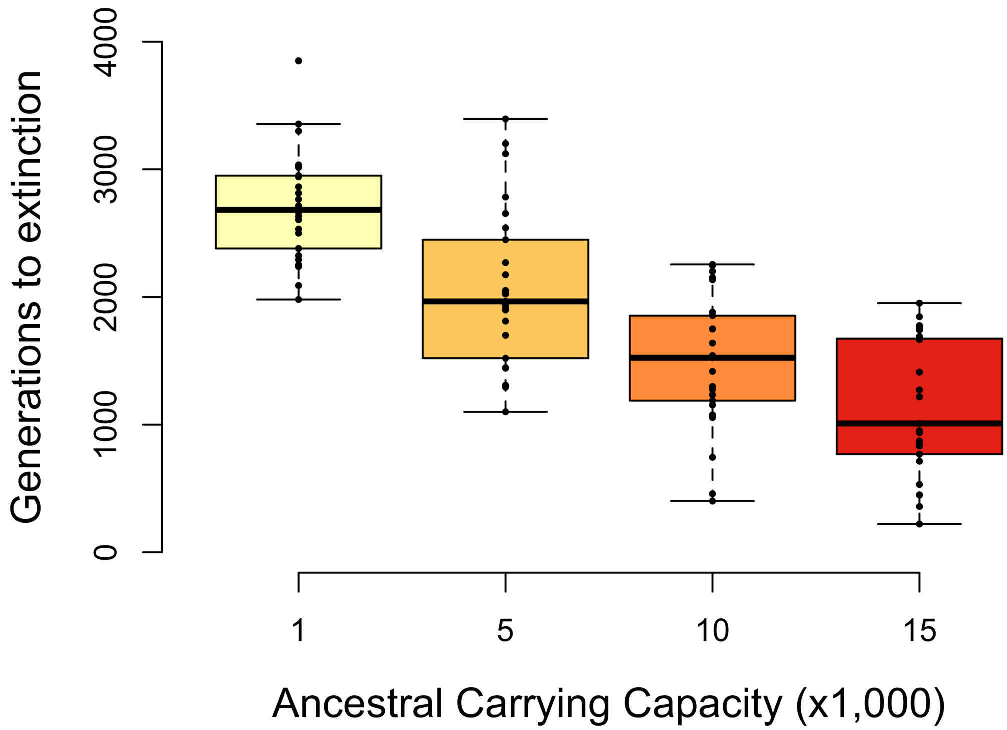


**Figure S7** Time to extinction following a population contraction under an *hmix* model of dominance when $K_{endangered}$=50. For each ancestral carrying capacity, 25 simulation replicates were run.

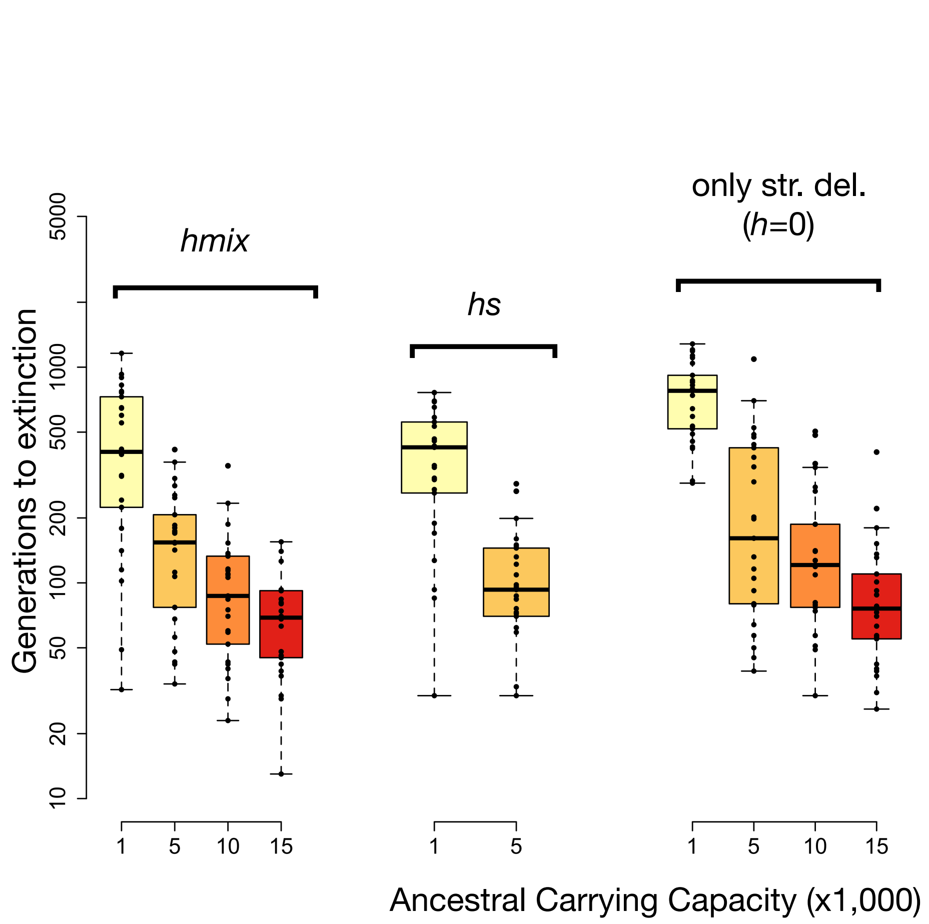


**Figure S8** Time to extinction following a contraction to $K_{endangered}$ = 25 comparing results under the full *hmix* and *hs* relationship models to those that only included the fraction of mutations in the *hmix* model that are strongly deleterious (*s* < -0.01 and *h*=0) or weakly/moderately deleterious (*s* $\geq$ -0.01 and *h*=0.25). Note that extinction times for strongly deleterious mutations are highly similar to the full models, despite these mutations making up only ~25% of new mutations under our assumed DFE (Kim et al. 2017). Also note that the y-axis is on a log scale. For parameter combination, 25 simulation replicates were run.

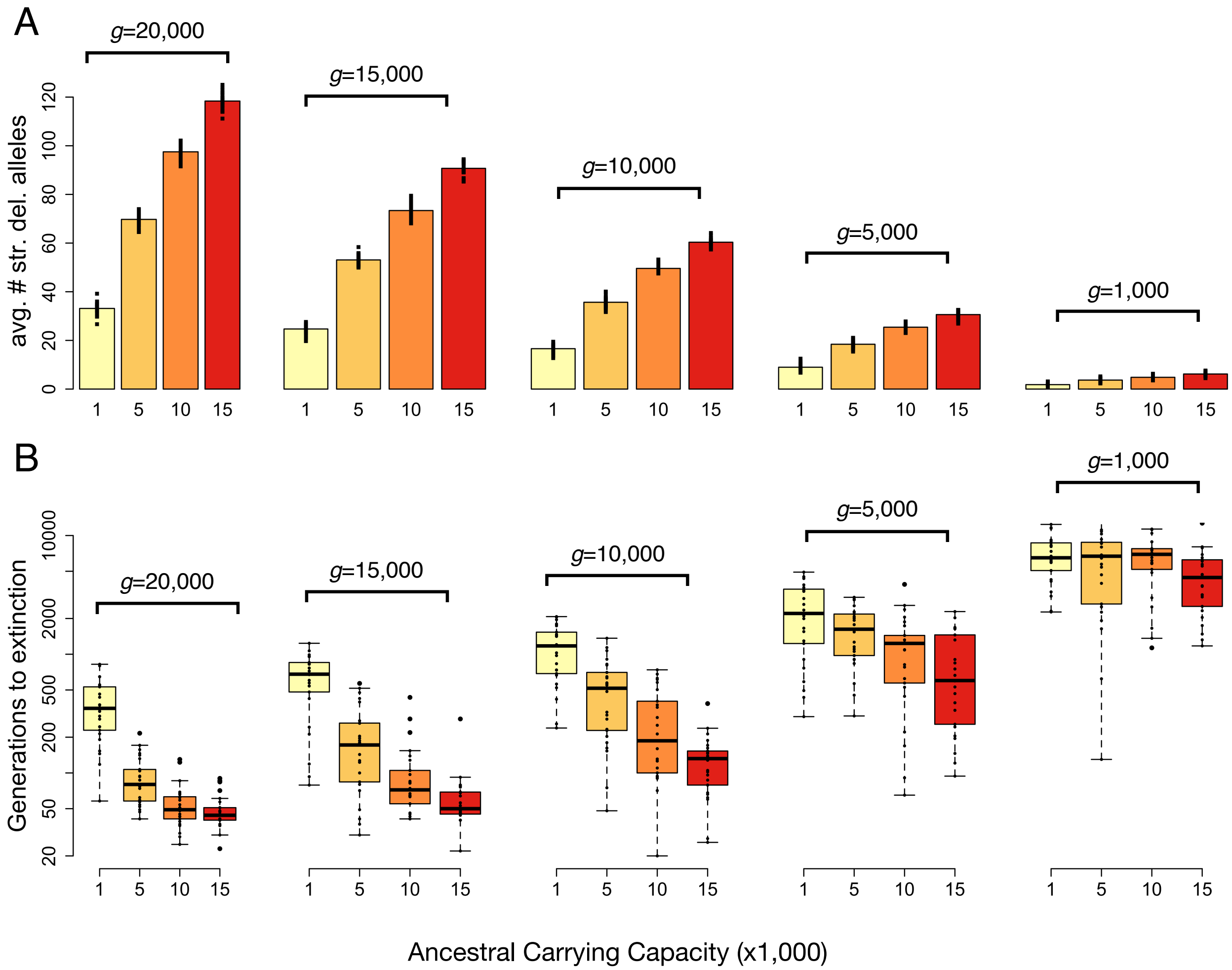


**Figure S9** Population contraction results when reducing the target size for deleterious mutations under a fully recessive model of dominance (*h*=0). (A) Average number of strongly deleterious alleles ($s < -0.01$) per individual in the ancestral populations prior to contraction. (B) Time to extinction following contraction from ancestral populations of varying size to an endangered population with K=25. Here *g* denotes the number of genes that can accumulate recessive deleterious mutations. Note that the y-axis is on a log scale. For parameter combination, 25 simulation replicates were run.

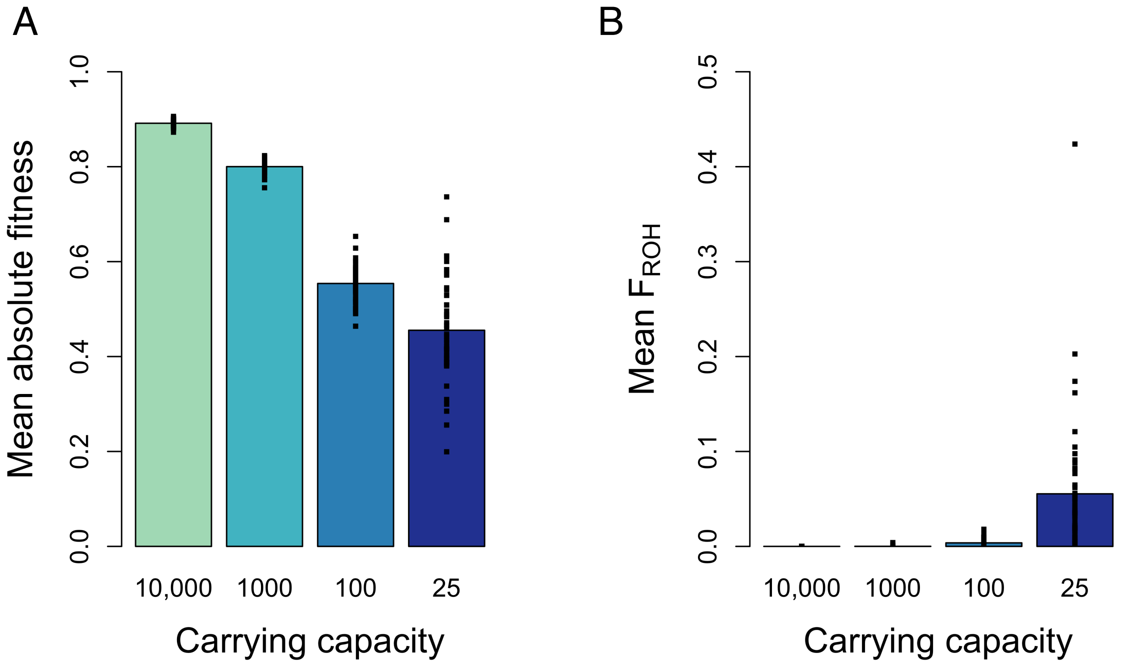


**Figure S10** Characteristics of the source populations used for genetic rescue. (A) Mean absolute fitness and (B) mean inbreeding coefficient ($F_{ROH}$) of the source population at the time of genetic rescue. For parameter combination, 50 simulation replicates were run.

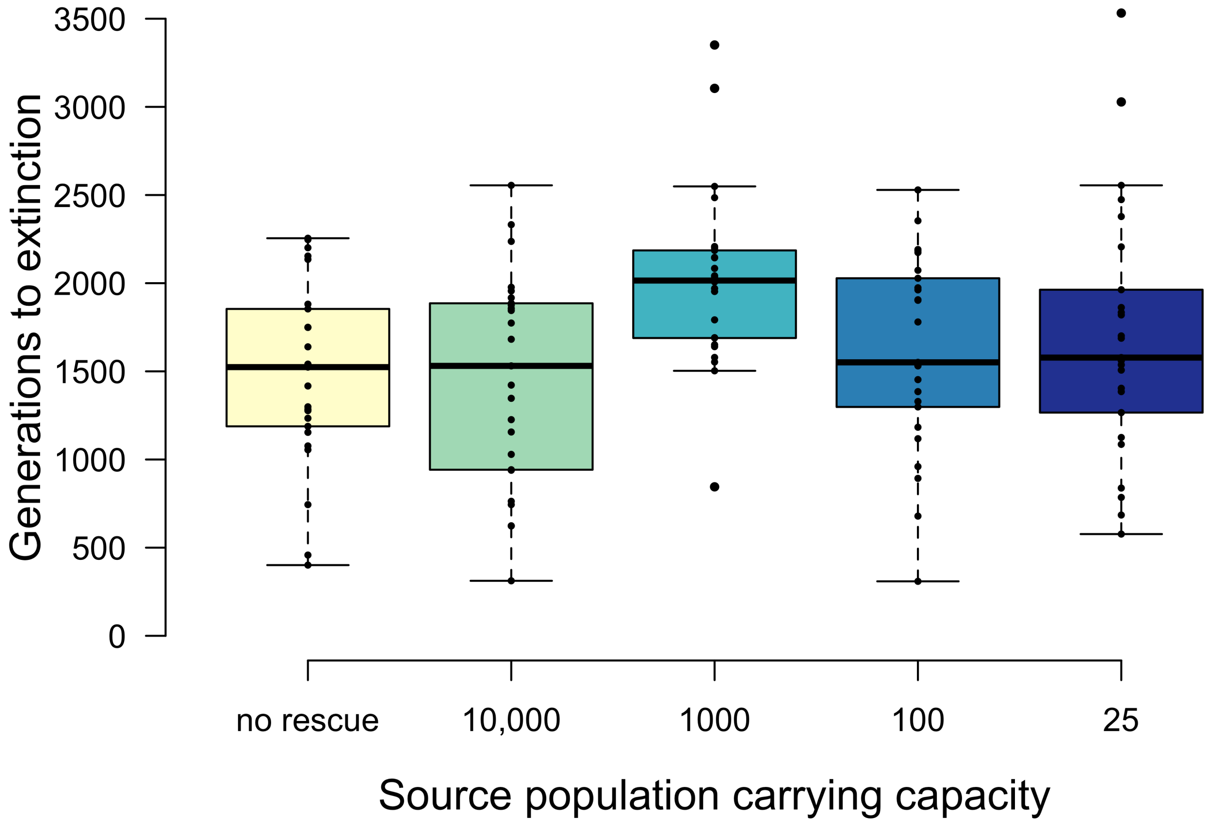


**Figure S11** Time to extinction following genetic rescue from source populations of varying size with recipient population carrying capacity of 50 under an *hmix* model of dominance. Rescue occurred when the recipient population decreased to 15 or fewer individuals. For each parameter setting, 25 simulation replicates were run.

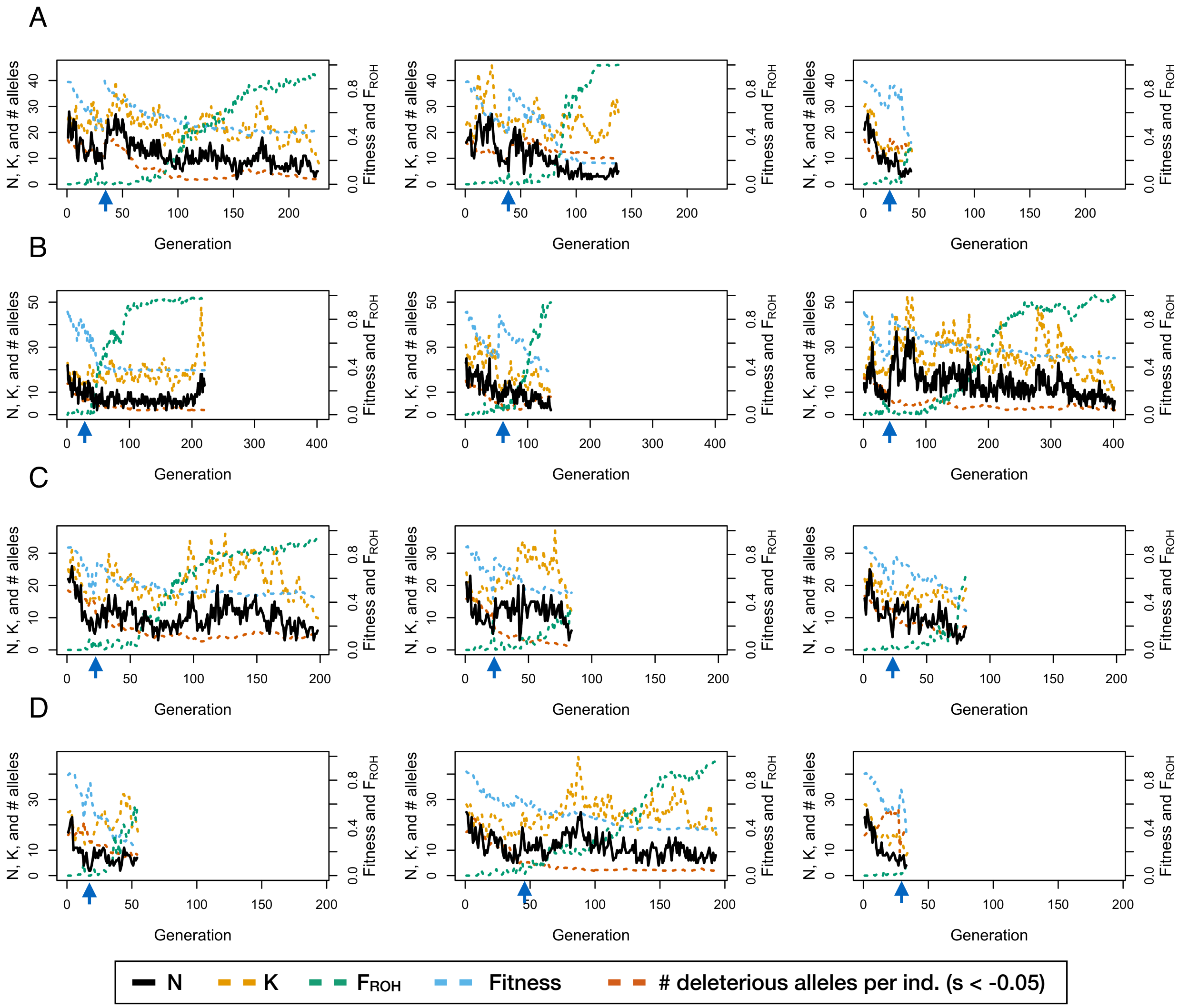


**Figure S12** Population trajectory of individual simulation replicates with $K_{endangered}$ = 25 and $K_{ancestral}$ = 10,000 following genetic rescue from source populations of varying demographic history: (A) $K_{source}$ = 10,000, (B) $K_{source}$ = 1,000 for 1,000 generations, (C) $K_{source}$ = 100 for 100 generations, and (D) $K_{source}$ = 25 for 10 generations. The timing of genetic rescue is indicated by the blue arrow. Three representative replicates are shown for each combination of parameters. Note that x and y axis scales differ for each set of three replicates.

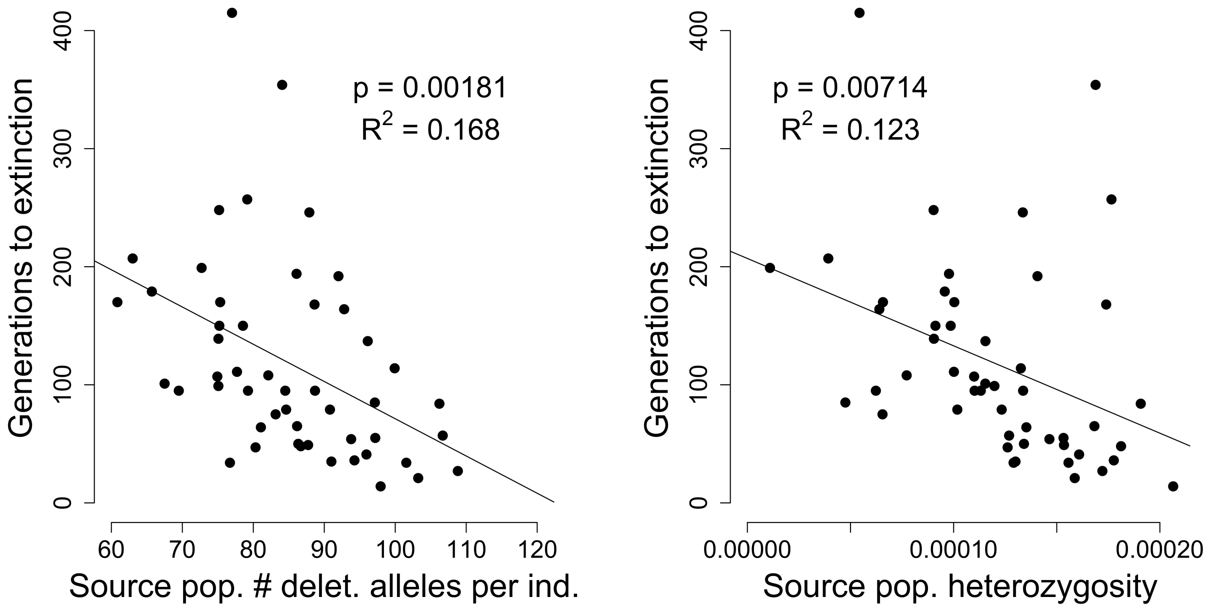


**Figure S13** Relationship between extinction times and source population deleterious variation and genetic diversity for the K=25 source population. (A) Extinction times are significantly negatively correlated with the average number of strongly deleterious alleles (*s* < -0.01) per individual in the source population at the time of translocation. (B) Extinction times are also significantly negatively correlated with average heterozygosity of the source population at the time of translocation.

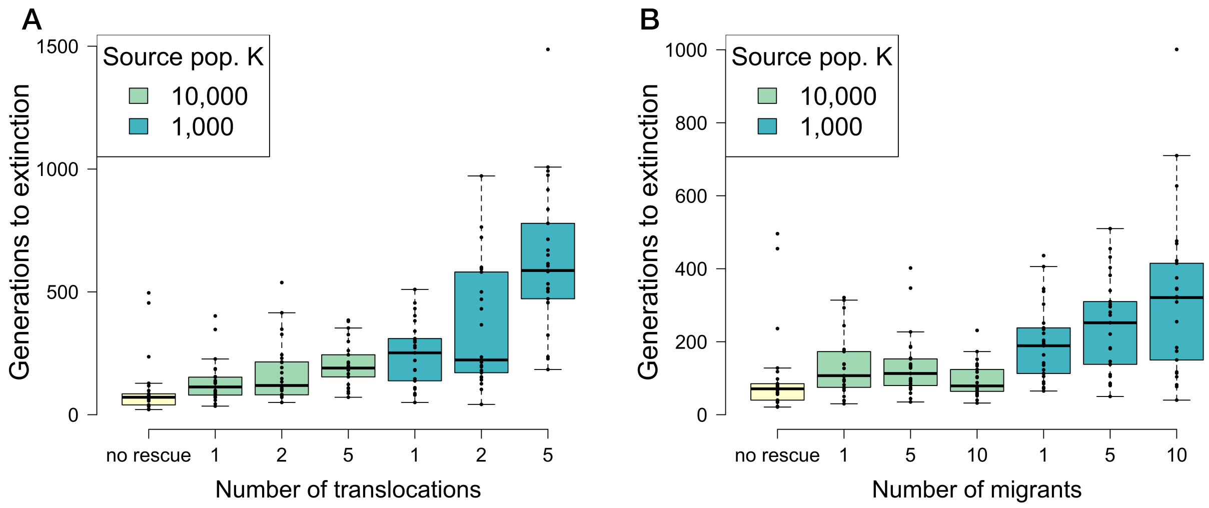


**Figure S14** Time to extinction following genetic rescue with varying numbers of translocation events and migrants with $K_{endangered} = 25$. (A) Results when varying the number of translocations from a moderate-sized (K=1,000) and large (K=10,000) source population, holding the number of migrants fixed at five. (B) Time to extinction when varying the number of migrants from a moderate-sized (K=1,000) and large (K=10,000) source population, holding the number of translocations fixed at one. For each parameter setting, 25 simulation replicates were run.
